# Modelling rheumatoid arthritis: A hybrid modelling framework to describe pannus formation in a small joint

**DOI:** 10.1101/2021.09.02.458714

**Authors:** Fiona R Macfarlane, Mark AJ Chaplain, Raluca Eftimie

## Abstract

Rheumatoid arthritis (RA) is a chronic inflammatory disorder that causes pain, swelling and stiffness in the joints, and negatively impacts the life of affected patients. The disease does not have a cure yet, as there are still many aspects of this complex disorder that are not fully understood. While mathematical models can shed light on some of these aspects, to date there are few such models that can be used to better understand the disease. As a first step in the mechanistic understanding of RA, in this study we introduce a new hybrid mathematical modelling framework that describes pannus formation in a small proximal interphalangeal (PIP) joint. We perform numerical simulations with this new model, to investigate the impact of different levels of immune cells (macrophages and fibroblasts) on the degradation of bone and cartilage. Since many model parameters are unknown and cannot be estimated due to a lack of experiments, we also perform a sensitivity analysis of model outputs to various model parameters (single parameters or combinations of parameters). Finally, we discuss how our model could be applied to investigate current treatments for RA, for example, methotrexate, TNF-inhibitors or tocilizumab, which can impact different model parameters.

## 1. Introduction

Rheumatoid arthritis (RA) is a chronic inflammatory disorder that affects over 1% of the worldwide population [1]. The characteristics of RA include persistent inflammation of joints, which contributes to the degradation of cartilage, and damage to the bone(s) within the joint [2, 3]. Along with symptoms related to inflammation within joints, RA can increase the risk of other health issues such as cardiovascular events [3, 4], reduced cognitive function in the brain, fibrotic disease in the lungs, osteoporosis and a greater risk of cancers [5]. The main symptoms of RA include inflammation, pain, swelling and stiffness of joints, fatigue and weight loss. Generally, the smaller joints in the hands and feet are most likely to be affected [4], for example, proximal interphalangeal (PIP) joints, which are the middle joints on the non-thumb fingers of the hand, are most commonly affected [6–9]. A key aspect of RA progression within a joint is the formation of a ‘pannus’ from the abnormally growing synovial membrane. The pannus is made up of mainly fibroblast-like synoviocytes (FLSs) and macrophage-like synoviocytes (MLSs). Through the production of inflammatory cytokines the proliferation, migration and cytokine secretion of these cell types increases leading to further inflammation. These cells also produce matrix degrading enzymes, like MMPs, which can breakdown cartilage and bone within a joint. In this work, we focus on these key features of pannus formation and growth and the subsequent cartilage and bone degradation. In the following subsection we provide further biological detail of these specific mechanisms. For more robust reviews of the biology of RA we direct the reader to the following papers [5, 10].

### 1.1. Key biological background

The synovial membrane is a soft connective tissue which lines synovial joints and allows for smooth movement through the secretion of lubricating synovial fluid [3, 10]. This well vascularised tissue consists of a intimal layer of evenly dispersed cells and a sub-lining comprised of extracellular matrix interspersed with collagen fibrils and other matrix proteins [11]. The porous structure of the synovial membrane allows for the diffusion of nutrients, oxygen and chemokines into the joint [12]. In a healthy joint, the intimal layer of the membrane is generally 1-2 cells thick and consists of fibroblast-like synoviocytes (FLSs) and macrophage-like synoviocytes (MLSs) evenly distributed and in equal amounts [11, 12]. Following the onset of RA, the synovial membrane expands through various inflammatory mechanisms, this growing membrane is known as a ‘pannus’ and can behave similarly to a locally invasive tumour spreading within the joint [11]. Through both an increase in the proliferation of FLSs and the infiltration of immune cells, such as bone-marrow derived macrophages, the synovial membrane can expand to around 10-20 cells in thickness [5, 11, 13–15]. In this work, we focus on resident macrophages, or macrophage-like synoviocytes, and resident fibroblasts, or fibroblast-like synoviocytes, which form the majority of the pannus. However, the pannus can also consist of other immune cell types, including leukocytes, plasma cells, T cells and mast cells. Pannus formation and growth is controlled through the expression of pro-inflammatory cytokines by the cells, promoting further inflammation of the synovial membrane. The cytokines involved are vast and varied, including those with roles in immune cell recruitment, immune cell activation, chemotaxis of cells and degradation of the cartilage/bone. There can be a high level of heterogeneity in macrophage origin and function in the context of RA [15], as well as FLS phenotypes and cytokine profiles [16]. In general, the pannus is a heterogeneous structure, consisting of diverse cellular and molecular signatures [11]. In recent years, distinct patterns have been recognised primarily according to composition, organisation and localisation of cellular infiltrates [11]. As a first approach, we consider homogenous cell populations whereby each cell within a population will exhibit the same phenotypic characteristics. However, we do implement stochasticity within the system through our modelling choices.

Cartilage is a connective tissue consisting of chondrocytes which produce a dense extra cellular matrix (ECM). Matrix metalloproteinases (MMPs) and tissue inhibitors of metalloproteinases (TIMPs) mediate cartilage destruction and are produced by B cells, FLSs and macrophages in the RA setting [3, 5, 10, 13, 17]. Cells within the pannus can also stimulate cartilage degradation via direct cell contact mechanisms [3, 14]. After cartilage has been damaged the bone underneath can become exposed. Bone erosion can be induced by cytokines that promote osteoclasts within the bone. In health, osteoclasts inhibit osteoblasts which produce new bone. In RA, the function of osteoclasts is increased, reducing the levels of new bone being formed leading to a reduction in bone formation [5, 10, 13]. As an initial formulation, for simplicity, we will consider both total cartilage and total bone densities rather than the individual components of each of these tissues.

Once initiated the outcomes of RA cannot be reversed, however disease progression can be slowed and the symptoms of the disease can be reduced through a variety of treatment approaches. There are several classes of RA treatment drugs including non-steroidal anti-inflammatory drugs (NSAIDs), steroids and disease modifying anti-rheumatic drugs (DMARDs). These drugs can be used alone or in conjunction with other treatment approaches. Biological DMARDs are generally used in conjunction with conventional DMARDs, increasing their efficacy, and are considered to have a strong benefit-to-risk profile [18, 19]. However, a large number of patients (approx. 40%) do not respond to the therapy, while others respond initially and then lose response over time [20]. Switching biologics is one approach considered for the patients with inadequate response to the initial treatment, although the second biologic might not be more effective than the first one [20]. For a full description of the drugs used in rheumatoid arthritis treatment we refer the reader to the following papers [4, 21, 22]. We discuss some specific drugs and the effects they have in the RA context in Section 3.4.

### 1.2. Previous mathematical descriptions of RA

Mathematical modelling is a useful tool to aid in the understanding of biological process at multiple spatial and temporal scales. In a recent review paper [23], we have considered existing mathematical models that aim to capture the mechanisms of rheumatoid arthritis in a variety of contexts. We refer the reader to the review paper for the full details, however we provide a short summary of the mathematical approaches reviewed here.

#### 1.2.1. Summary of models reviewed in [23]

Systems of ordinary differential equations (ODEs), which are non-spatial deterministic continuous equations that describe the time evolution of a variable of interest, are the most common modelling approach used to describe the evolution of RA. Single compartment ODE models have been used to model joint erosion [24], the interactions between generic pro- and anti-inflammatory cytokines [25], the role of the cytokine TNF-*α* [26, 27], the interactions of immune cells within the RA environment and the drug Tocilizumab [28]. More complex, multi-compartment ODE approaches have also been used to consider the circadian dynamics involved in the progression of rheumatoid arthritis [29] and the inflammatory and invasive processes occurring at the cartilage-pannus interface [30]. Furthermore, a number of ODE based approaches consider the pharmacokinetics and pharmacodynamics of various drugs used to treat RA [31–38].

To account for spatio-temporal features of RA, systems of partial differential equations (PDEs) have been utilised. PDEs are deterministic continuous equations that can describe the spatial and temporal evolution of a variable of interest. PDEs are less commonly used to describe RA specific processes, in comparison to ODEs. For example, Moise *et. al*. [39] consider a three-compartment model to describe the spatio-temporal interactions between immune cells, cytokines and drugs in the synovial membrane, synovial fluid and cartilage of a joint.

Deterministic methods such as ODEs and PDEs cannot capture the potential variability or stochasticity within a biological system. Stochastic mathematical and computational models describe the interactions between the different components of the system, or the transitions between different states of these components as probabilistic. Such models have been mainly applied in the context of treatment decisions [40], to analyse RA incidence rates [41], or to predict radiological progression within RA [42, 43]. Moreover, a subset of stochastic Markov chain models have focused on assessing the cost-effectiveness of single or combined RA treatments [44, 44–47].

#### 1.2.2. Further stochastic and hybrid (deterministic-stochastic) modelling approaches

The few models existent in the literature are mainly deterministic and are given by differential equations. However it has been suggested that accounting for stochasticity within RA may be key in understanding the evolution of the disease [10, 48] and predicting the success of RA treatments [49]. Along with the stochastic models described in Section 1.2.1, a small number of studies have considered Boolean network models for RA [50, 51].

A different approach to incorporate stochasticity at a cellular level within mathematical models is to use an individual-based (IB) (or agent-based) modelling approach [52, 53]. This approach allows each cell to be described as an individual agent which follows a set of predetermined rules. These agents reside in a defined spatial domain. This domain can be constrained as a lattice, where agents are restricted to moving between lattice positions, or more realistically an off-lattice approach can be utilised where agents have freedom in direction of movement. Off-lattice individual-based modelling has been used in multiple areas of mathematical biology research including modelling the cellular immune response to viruses [54] and cancer [55]. Individual-based models can be computationally expensive especially when modelling a large number of agents. This expense can be reduced by using a hybrid modelling approach. For example, when modelling a chemical we may not necessarily be interested in modelling each individual molecule but the total local concentration instead. To model these more continuous aspects a deterministic approach can be used, where the total local concentration is described rather than each molecule, this can be viewed as a tissue-level approach. To allow us to consider both the stochastic cellular level scale and deterministic tissue level dynamics of the system we can use a hybrid multiscale modelling approach [56]. To model the deterministic components we can use classical methods such as differential equations. Hybrid approaches have been used to model various biological phenomena such as monocyte migration in the vasculature [57], the cellular immune response to sepsis [58], inflammation in chronic obstructive pulmonary disease [59] or various mechanisms of cancer growth and development [60–63]. In a similar way, in the context of rheumatoid arthritis, using multi-scale hybrid modelling approaches may be valuable in modelling disease progression and predicting the success of RA treatment.

These hybrid modelling approaches allow for the dynamics of single cells within a system to be investigated while chemical concentrations and tissue-level effects are modelled deterministically. This is useful in the context of RA as it has been found that there exists high levels of phenotypic and spatial heterogeneity of cells within the synovial membrane [64]. Moreover, understanding these single cell phenomena further has been highlighted as an important step in the identification of treatment targets [65–67]. To investigate these single cell dynamics, we propose a hybrid (stochastic-discrete) modelling method would be appropriate to describe the cellular dynamics in the progression of RA.

### 1.3. Overview of paper

Building upon previous on-lattice hybrid models to describe interacting cell and chemical populations [68, 69], we develop an off-lattice hybrid model as a first step towards building a more realistic model of arthritic destruction within a small PIP joint. More specifically, we incorporate the dynamics of two resident cell populations, fibroblasts (or fibroblast-like synoviocytes) and macrophages (or macrophage-like synoviocytes) through a stochastic off-lattice individual-based modelling approach. The dynamics of cartilage and bone density, along with the evolution of MMPs are described via a deterministic PDE approach. In the model we consider the simplified interactions of these components which can lead to destruction of cartilage and subsequently the bone within the joint. The methods used to implement each of the mechanisms described are those which could be used to include the necessary biological detail as the framework is improved upon in future iterations of the model. The paper is structured as follows. In Section 2, we describe the details of the framework and how each biological mechanism is described mathematically. In Section 3, as an example we highlight some initial results of the framework and perform a sensitivity analysis to investigate the role of the parameters in the RA system. We additionally describe how the framework could be related to current rheumatoid arthritis treatments. Finally in Section 4 we discuss the key outputs of our framework and their biological relevance, the requirements to validate the framework and the plans for developing the framework further.

## 2. The discrete model

In this section we introduce the hybrid modelling framework used to describe pannus formation in a joint affected by rheumatoid arthritis. In the framework we consider 5 components: bone and cartilage densities, resident fibroblasts (or fibroblast-like synoviocytes) and resident macrophages (or macrophage-like synoviocytes) and the degrading proteases, matrix metalloproteinases (MMPs). Note, we include only fibroblasts and macrophages as they contribute to the majority of the pannus, however further cell types could be considered in future iterations of the modelling framework. We use an off-lattice individual-based modelling approach to describe the dynamics of the cells, coupled with discretised PDEs to model cartilage density, bone density and MMP concentration. In Section 2.1, we describe the set-up of the initial spatial domain to replicate a PIP joint. The mathematical methods used to describe both fibroblast and macrophage dynamics are presented in Section 2.2. Section 2.3 contains the mechanisms used to describe the evolution of MMPs in the system. The methods used to describe both cartilage and bone density are given in Section 2.4. Finally, we describe how the on and off-lattice components of the model are coupled in Section 2.5.

### 2.1. Set-up of the domain

We consider a 2D spatial domain with *x* ∈ [*x_l_*, *x_h_*] and *y* ∈ [*y_l_, y_h_*]. A schematic diagram of the spatial domain and initial condition is provided in Figure 1. As described in Section 1.1, small joints in the hand are most commonly affected by rheumatoid arthritis and can be affected in early stages of the disease. Therefore we formulate the model in the context of rheumatoid arthritis within a small PIP joint. There would be no difficulty in reframing the model in the context of other RA infected joints. To replicate a small joint we consider the joint space to contain two bone ends, surrounded by cartilage with a small space between the two bones. As a simplification, in the model we consider the bone and cartilage domains to be rectangular. In healthy joints, the synovial membrane is around 1-2 cells thick consisting of macrophages and fibroblasts which are evenly distributed and in equal amounts [11, 12]. Therefore, we position macrophages and fibroblasts randomly in two layers at the boundaries of the domain, where the width of the membrane is approximately 2 cells thick. To define the spatial domain, we consider the width, length or height of each of these areas by defining the following parameters *W*, the descriptions of these parameters are available in Table 1.

**Figure 1:**
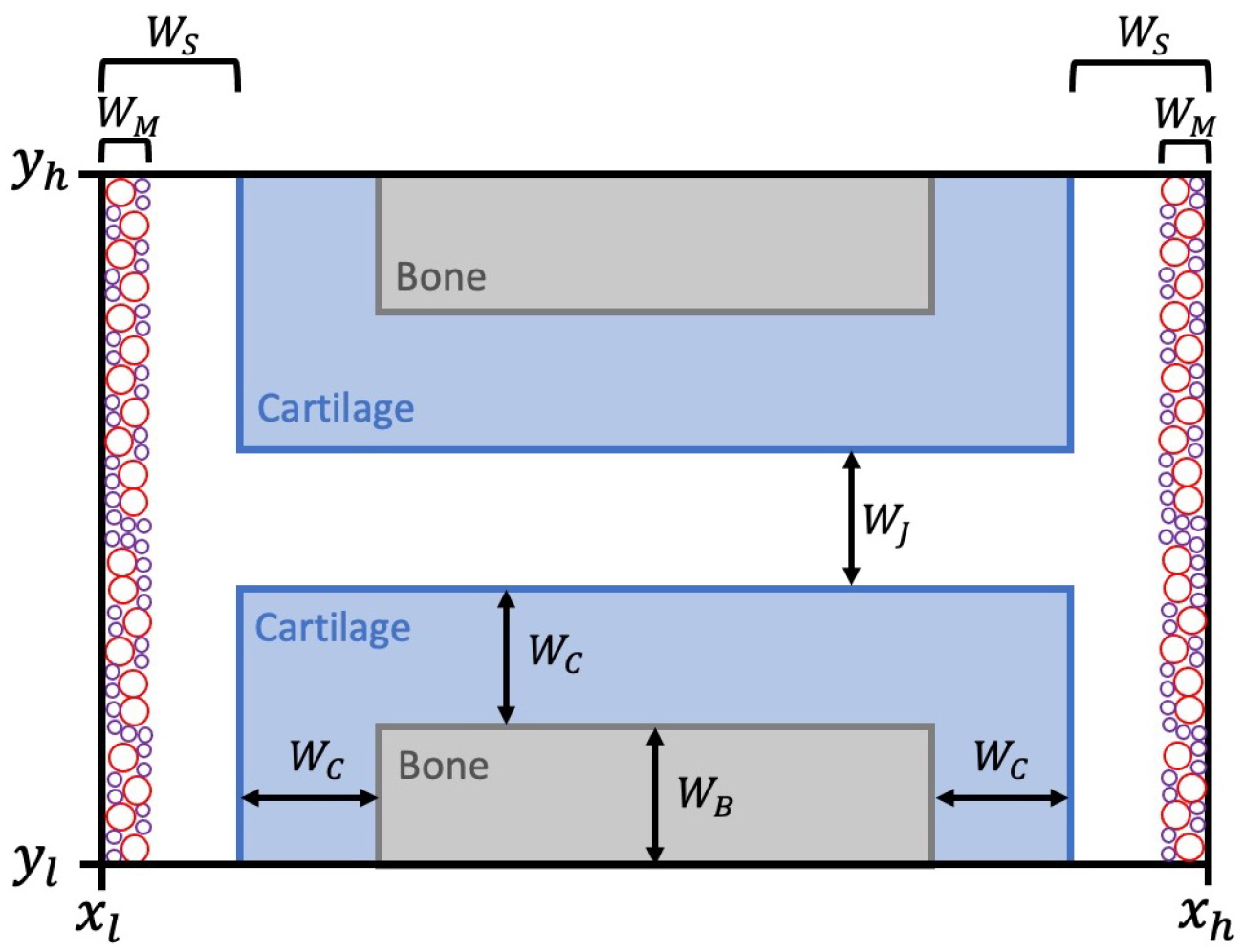
The initial spatial domain of the hybrid model. Shaded grey area represents the areas where bone is initially present, while shaded blue represents the areas where cartilage is initially present. The red and purple circles represent macrophages and fibroblasts, respectively. Note, that the initial position of both cell types of cells is randomised, so the positions here are just an example. The white space represents synovial fluid in which cells can freely move and proliferate, *e.g*., essentially free space in the system. Each of the parameters *W* are shown on the diagram represent the length, height or width of each of the areas within the domain. Note, the diagram is not to scale.

**Table 1:** Descriptions of the parameters used to set-up the initial spatial domain of the hybrid model as shown in Figure 1.

| Symbol | Description |
| --- | --- |
| $W_B$ | the height of bone protruding into the domain |
| $W_C$ | the width of the cartilage that surrounds the bone |
| $W_J$ | the width of the joint space between the two bone ends |
| $W_M$ | the initial width of the synovial membrane |
| $W_S$ | the space between the cartilage and the edge of the domain |

### 2.2. Cell dynamics

Fibroblasts and macrophage populations are included in the model using an off-lattice individual-based modelling approach. We track the total number of fibroblasts and macrophages over time, denoted by *N_F_*(*t*) and *N_M_*(*t*), respectively. To take into account the physical size of the cells, each cell within both populations is tracked by the spatial position of the cell centre and the radius of the cell. To allow for simplicity we consider all cells to be perfectly spherical (*i.e*. the radius of the cell is consistent) and we further consider that the cell size does not change over time. We consider homogeneous populations of fibroblasts and macrophages, that have the radii *R_F_* for the fibroblasts and *R_M_* for the macrophages.

Initially, we randomly place *N_F_* (0) fibroblasts and *N_M_* (0) macrophages in the areas denoted by width *W_M_*, as shown in Figure 1. To ensure physical space is taken into account, for each cell a desired initial position is chosen then checked for overlap with previously placed cells, cartilage, bone or the domain boundary as we consider a volume-exclusion process, whereby only one cell can occupy a particular area of free space. If there is overlap, a new position is chosen for that cell and the process repeats, until all cells are placed.

To replicate growth and invasion of the pannus into the joint space, we allow both cell types to undergo mechanisms of random motion/migration and proliferation within the individual-based model, as described in the following subsections. For simplicity, we consider both the fibroblast and macrophage populations to be homogeneous, that is each fibroblast will have the same potential to divide, die or migrate and similarly, each macrophage will have the same potential to divide, die or migrate. Furthermore, we impose zero-flux boundary conditions ensuring all cells remain within the spatial domain. That is, we consider only cells that reside within the simulated joint and omit the influx or recruitment of external immune cells into the domain.

#### Cell movement

We incorporate random cell movement for both fibroblasts and macrophages using the same mechanism and allow different probabilities of movement for each cell type. At every time-step each cell can move randomly with probability, λ_*F*_ or λ_*M*_, if a fibroblast or macrophage, respectively. If the cell is permitted to move, then a new direction is chosen from an angle *θ* between 0 and 2*π* for the cell to move in, as shown in Figure 2. The new position is calculated as a jump of length Δ_*x*_, to ensure proper scaling of movement probabilities. The movement to this desired position is only permitted if the new position is within the boundary, does not overlap with any other cells or with cartilage/bone density. If any of these properties prevent movement, the movement jump is aborted.

**Figure 2:**
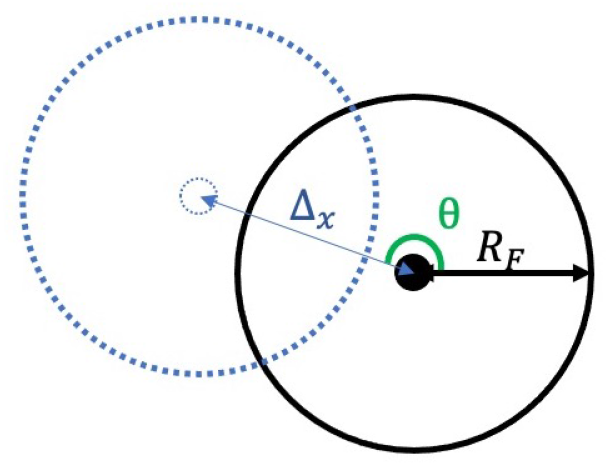
Schematic of the off-lattice cell movement mechanism. The solid black circle represents the initial cell position where the cell has radius *R*. A desired position is chosen by selecting a random angle *θ* ∈ [0, 2π] and then placing the cell centre at a distance of Δ_*x*_ from the original position in the direction *θ*. The desired position (blue dashed circle) is chosen as the cell’s updated position if no other cells, cartilage or bone overlap this spatial position. If this position does overlap the cell movement process is aborted.

#### Cell proliferation

We incorporate cell division and death for both fibroblasts and macrophages using the same mechanism and allow different probabilities of division and death for each cell type. It has been shown that a major factor controlling cell division in fibroblasts is a random event controlled by a transition probability [70]. Therefore, to incorporate cell division into our model we allow for this random event by allocating a probability of division at each time-step to each cell. This random event occurs once the cell has reached a minimum size, however for simplicity in this model we do not incorporate cell cycle or cell size, these details could be explicitly considered in future developments of the model. At every time-step, each cell has a probability of dividing given by *α_F_* or *α_M_*, for fibroblasts or macrophages respectively. If the cell is permitted to divide, then the cell splits into two identical daughter cells. One is placed on the parent cell’s original position and the other is placed in a chosen position adjacent to this, as shown in Figure 3. This adjacent position is chosen from an angle *θ* between 0 and 2*π*. The new position is set at a length of twice the radius of the cell *R_F_* or *R_M_*, *i.e*., the cell’s diameter. This new cell is only placed on this position if the position is within the boundary, does not overlap with any other cells or with cartilage/bone density. If any of these properties prevent division, the process is aborted and the parent cell remains in the original position.

**Figure 3:**
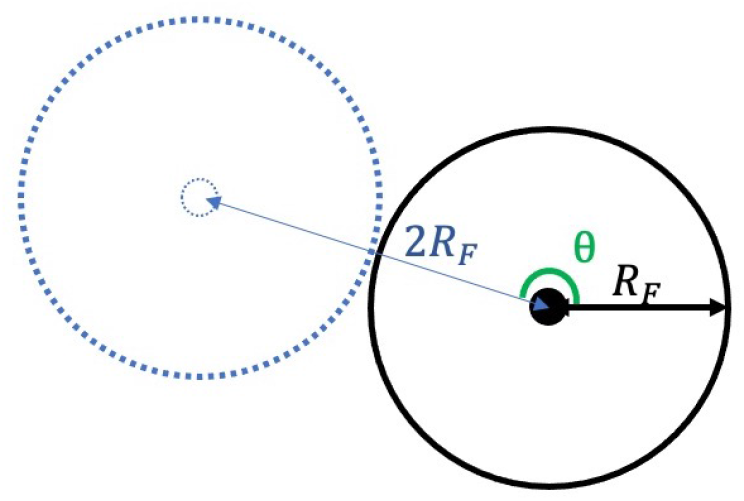
Schematic of the off-lattice cell division mechanism. The solid black circle represents the original cell position where the cell has radius *R*. The cell will divide into two identical daughter cells, one cell will remain on the original cell’s position and the other will be placed at a desired position adjacent to the original cell’s position. A desired position for the second daughter cell is chosen by selecting a random angle *θ* ∈ [0, 2*π*] and then placing the cell centre at a distance of 2R, *i.e*., the cell’s diameter, from the original position in the direction *θ*. The desired position (blue dashed circle) is chosen as the daughter cell’s position if no other cells, cartilage or bone overlap this spatial position. If this position does overlap the cell division process is aborted and the original cell remains.

Furthermore, at every time-step, each cell has a probability of dying. The probability of dying is given by *κ_F_* or *κ_M_*, for fibroblasts or macrophages respectively. If the cell is assigned to die at a given time-step, it is simply removed from the simulation instantly.

### 2.3. MMP dynamics

Since matrix metalloproteinases (MMPs) produced by fibroblasts and macrophages mediate cartilage destruction and bone destruction we include the dynamics of MMPs in to the mathematical model. We describe the dynamics of MMPs in the system as a deterministic process, as we are interested in the total local concentration of MMPs rather than each individual molecule. We use a similar approach to Moise *et. al*. [39] and include the diffusion, production and decay of MMPs. As MMPs are secreted by both fibroblasts and macrophages, we consider the density of cells to contribute to the local MMP concentration. In the RA setting, tissue inhibitors of metalloproteinases (TIMPs) regulate the activity of MMPs, however for simplicity, we take into account the actions of TIMPs implicitly by including a decay rate of MMPs and imposing a local limit on the concentration of MMPs at any spatial position to ensure the local concentration is bounded. Mathematically these assumptions can be captured by the PDE,

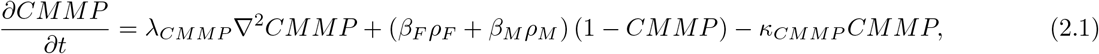

where *CMMP* = *CMMP*(**x**, *t*) is the local concentration of MMPs at position **x** at time *t*, λ*_CMMP_* is the diffusivity of MMPs, *β_F_* and *β_M_* are the rates of production of MMPs by fibroblasts and macrophages, respectively, *ρ_F_* = *ρ_F_* (**x**, *t*) and *ρ*_*M*_ = *ρ*_*M*_(**x**, *t*) are the cell densities of fibroblasts and macrophages, respectively, and *κ_CMMP_* is the decay rate of MMPs. We solve the PDE with zero-Neumann boundary conditions to allow all MMPs to remain within the domain. We note that as MMPs are a small molecule, MMPs can enter the areas of cartilage and bone density freely and there are no spatial restrictions on the MMPs diffusion within the domain.

### 2.4. Cartilage and Bone dynamics

In the model, we are interested in the destruction of cartilage and bone by MMPs. Therefore, we consider the density of cartilage and bone within the spatial domain over time. For simplicity, we do not include the biological components of cartilage or bone, and consider these tissues as a whole. We model the evolution of cartilage and bone through the PDEs,

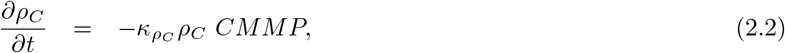

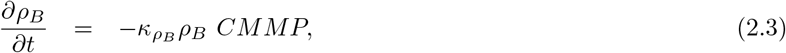

where *ρ_C_* = *ρ_C_* (**x**, *t*) and *ρ_B_* = *ρ_B_* (**x**, *t*) are the densities of cartilage and bone, respectively. Here, *κ_ρ_C__* and *κ_Pρ_B__* are the rates of degradation of cartilage and bone by MMPs, *CMMP* = *CMMP*(**x**, *t*), respectively. We solve these equations with zero-Neumann boundary conditions to ensure all components remain within the spatial domain. As an initial formulation of the model, we do not allow for the growth of cartilage or bone which could be incorporated by adding more levels of complexity.

### 2.5. Linking the individual-based and PDE components of the model

As the modelling framework utilises a hybrid approach (*i.e*. a stochastic off-lattice model for cell dynamics and deterministic PDEs for MMPs, cartilage and bone evolution) we have to ensure that the two approaches allow for interactions between the stochastic and deterministic components. For numerical simulations of the model we discretise Equations (2.1)–(2.3) such that each of these densities is defined on a discrete lattice, the discretisation method is provided in Appendix A. The description of how the off-lattice components are then coupled with the discrete lattice components are described in the following paragraphs.

#### Secretion of MMPs

In Equation (2.1) at the tissue-level scale, the production of MMPs is dependent on the density of fibroblasts and macrophages. At the cellular level scale, this translates to the secretion of MMPs by each of the cells. For simplicity, we assume that cells produce MMPs at the centre position of the cell. To allow the concentration of MMPs to remain on our discretised grid, we calculate the nearest grid-position to the cell centre, and add the desired MMP concentration to this grid position.

#### Volume-exclusion

To ensure that the cells do not move or divide onto areas of the domain that contains bone or cartilage we need to evaluate the position of the whole cell on the grid. If no cartilage or bone degradation has occurred we simply check whether the new desired position is within ±*R* (radius) of the initial areas of the domain containing cartilage or bone, as depicted in Figure 1. That is, we check any part of the cell will be within the initial cartilage or bone domain areas. If bone or cartilage degradation has occurred then we want to ensure that the cell can now move into this free space. To do this we find equally distributed points on each cell’s circumference and then calculate the nearest grid-position of each of these points. We then check whether there is cartilage or bone present on those positions, to ensure the cell is able to move. Note, here we make the assumption that a cell can only inhabit an area of space if the cartilage and bone density is zero, however in future work we could allow cells to inhabit areas if these densities are below some threshold value, if appropriate.

## 3. Numerical simulations and sensitivity analysis

To visualise the hybrid modelling framework developed in Section 2 we run numerical simulations using MATLAB, the details of the explicit method used to solve the PDEs is given in Appendix A. Every iteration of the simulation is run with the same initial condition. Each simulation is run over a 20 day time-frame as differences in the outcome of the model between runs with different parameter set-ups could be observed over this time-frame. A schematic overview of the key components, biological mechanisms and key parameters in the model is provided in Figure 4. To run these simulations we have to choose values for the parameters within the model, we display these parameters in Table 2. We then provide some example results of the model focusing on the aggressiveness of pannus growth through varying the division probabilities of macrophages and fibroblasts in Subsection 3.2. To investigate the role of each parameter within the model we perform a one-at-a-time sensitivity analysis in Subsection 3.3. Different therapeutic treatments can target specific mechanisms within the formation of the pannus and destruction of the joint, implicitly we can relate these therapeutics to varying single parameters within the model or varying combinations of parameters, we discuss this in Subsection 3.4.

**Figure 4:**
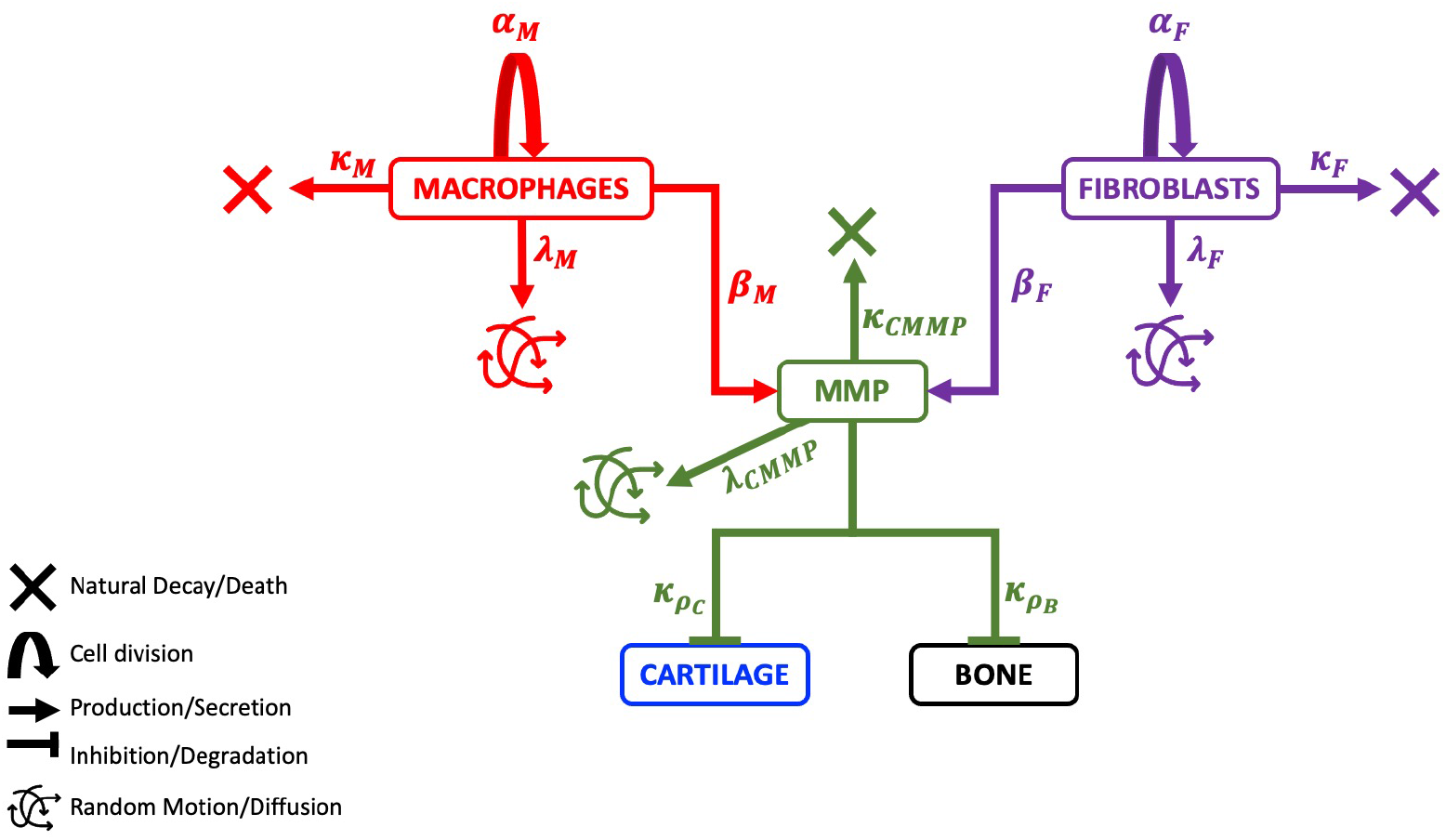
Schematic describing the components, mechanisms and parameters included in the hybrid model. Here, *α_F_* and *α_M_* represent the probability of cell division for fibroblasts and macrophages, respectively, while *κ_F_* and *κ_M_*, represent the probability of cell death. The probability of cell movement for fibroblasts and macrophages is given by λ*_F_* and λ*_M_*, respectively. The MMP secretion rates of fibroblasts and macrophages are given by *β_F_* and *β_M_*, respectively. The MMPs decay naturally at the rate *κ_CMMP_* and can diffuse in the spatial domain at the rate λ*_CMMP_*. The MMPs additionally degrade both cartilage and bone at the rates *κ_ρ_C__* and *κ_ρ_B__*, respectively.

**Table 2:** The parameters used in numerical simulations of the model. Unless a reference is given the parameters are estimated or calculated. Descriptions of how each value is chosen are provided in Appendix C.

| Symbol | Description | Value |
| --- | --- | --- |
| $\Delta_{tchem}$ | time-step length for deterministic components | $1 \times 10^{-4}$ days |
| $\Delta_{tcells}$ | time-step length for stochastic components | $1 \times 10^{-2}$ days |
| $\Delta_x$ | grid-step length in $x$ direction | $1 \times 10^{-3}$ cm |
| $\Delta_y$ | grid-step length in $y$ direction | $1 \times 10^{-3}$ cm |
| $W_B$ | initial thickness of bone | 0.1 cm |
| $W_C$ | initial thickness of cartilage | 0.04 cm [7] |
| $W_J$ | initial width of joint space | 0.01 cm [6, 8] |
| $W_M$ | initial width of synovial membrane | 0.004 cm [11] |
| $W_S$ | initial width between edge of domain and the cartilage | $2W_M$ cm |
| $x_l$ | minimum $x$ value of domain | 0 cm |
| $x_h$ | maximum $x$ value of domain | 0.3 cm |
| $y_l$ | minimum $y$ value of domain | 0 cm |
| $y_h$ | maximum $y$ value of domain | $W_J + 2W_C + 2W_B$ cm |
| $\alpha_F$ | probability of a fibroblast dividing | $0.33\Delta_{tcells}$ [39, 71] |
| $\alpha_M$ | probability of a macrophage dividing | $0.33\Delta_{tcells}$ |
| $\beta_F$ | concentration of MMPs produced by fibroblasts | $13.912 \times 10^{-5}\Delta_{tcells}$ [39] |
| $\beta_M$ | concentration of MMPs produced by macrophages | $5.2717 \times 10^{-5}\Delta_{tcells}$ [39] |
| $\kappa_F$ | probability of a fibroblast undergoing apoptosis | $0.03\Delta_{tcells}$ [39, 72] |
| $\kappa_M$ | probability of a macrophage undergoing apoptosis | $0.033\Delta_{tcells}$ [39, 73] |
| $\lambda_F$ | probability of a fibroblast moving via undirected motion | $\frac{\Delta_{tcells}}{\Delta_x^2} 8.64 \times 10^{-7}$ [39, 74] |
| $\lambda_M$ | probability of a macrophage moving via undirected motion | $\frac{\Delta_{tcells}}{\Delta_x^2} 8.64 \times 10^{-7}$ [39, 74] |
| $N_F(0)$ | initial number of fibroblasts | 200 cells. |
| $N_M(0)$ | initial number of macrophages | 200 cells. |
| $R_F$ | radius of a single fibroblast | $6.5 \times 10^{-4}$ cm [75]. |
| $R_M$ | radius of a single macrophage | $10.5 \times 10^{-4}$ cm [76] |
| $\lambda_{CMMP}$ | diffusivity of MMPs | $6.59 \times 10^{-2}$ cm<sup>2</sup> day<sup>-1</sup> [39, 74] |
| $\kappa_{CMMP}$ | natural decay rate of MMPs | 0.138 day<sup>-1</sup> [39, 77] |
| $\kappa_{\rho_C}$ | decay rate of cartilage by MMPs | $\frac{4.44 \times 10^3}{1.15} \rho_C(0)$ day<sup>-1</sup> [39] |
| $\kappa_{\rho_B}$ | decay rate of bone by MMPs | 0.1 $\kappa_{\rho_C}$ |

### 3.1. Parameterising the model

To parameterise the model, where possible, we use data from the literature and previous mathematical models. The parameters used in the numerical simulations of the model, unless stated otherwise are those shown in Table 2. We provide in Appendix C a full description of the choices of these parameters and how they are calculated.

### 3.2. Increasing the division probabilities of macrophages and fibroblasts

As an example of the simulation results of the model we consider the cases where all parameter values are taken to be the values given in Table 2. The simulation results for the first 20 days are shown in Figure 5. We plot the spatial distributions of cells, bone and cartilage at four time-points in the left-hand panels. We additionally plot the average number of fibroblasts, average number of macrophages, average global MMP concentration, average global cartilage density and average global bone density from 5 runs of the simulation in the right-hand plots. Here, global concentration/density refers to the sum of the MMP concentration or cartilage/bone densities across the whole spatial domain. We show the standard deviation between the 5 runs as the shaded area on each plot. We can see from these results that in this parameter setting, the cartilage density does decrease as MMP concentration increases, however no bone degradation occurs. In all of the line plots the standard deviation between runs is relatively low, especially in MMP concentration and the cartilage density plots.

**Figure 5:**
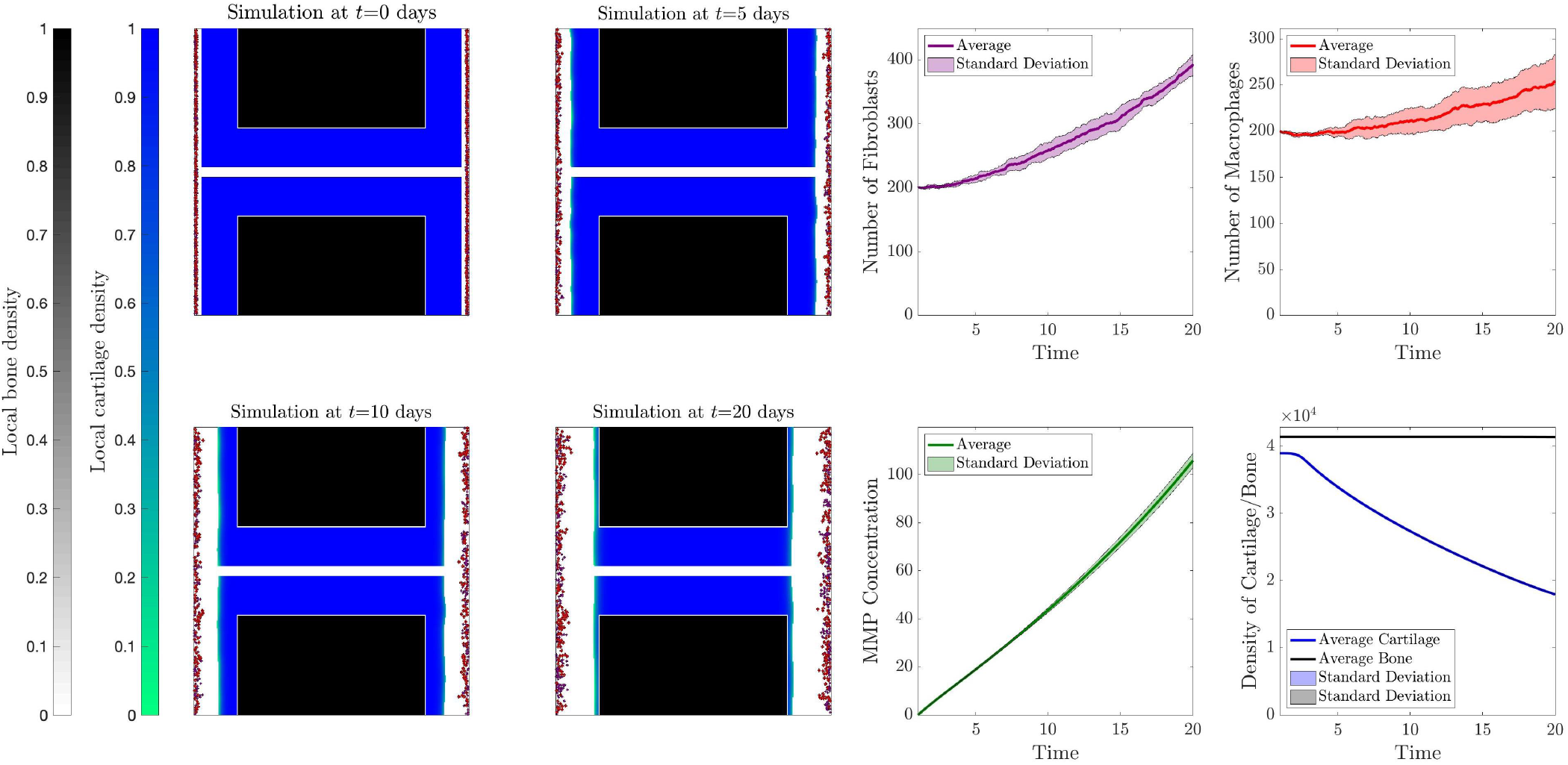
Example results where all parameters are those given in Table 2. **Left-hand plots:** Panels show the visualisation of spatial results of one of the runs of the simulation at the time-points *t*={0, 5, 10, 20} days. The red dots are macrophages, the purple dots are fibroblasts, the blue-green surface is cartilage density and the black-grey surface is bone density. White space represents space in which cells can move freely in the joint, *e.g*., synovial fluid. **Right-hand plots:** The plots show the cell number, concentration or density over time averaged over 5 simulation runs with the standard deviation shaded. The number of fibroblasts is given in purple, the number of macrophages in red, the global MMP concentration in green, the global cartilage density in blue and the global bone density in black.

We can also consider situations to replicate a biologically motivated scenario where immune activity has been amplified, as we would expect in more aggressive forms of rheumatoid arthritis. In Figure 6 we show results of the model where immune cell proliferation is increased ten-fold. From these results, we observe a larger increase in the number of fibroblasts and macrophages over time as expected. We also see that the concentration of MMPs is increased in comparison to the results shown in Figure 5. In this case, the cartilage degradation is more severe than the previous case, however the bone density is retained. Once again, we observe low standard deviation between runs for all of the model outputs.

**Figure 6:**
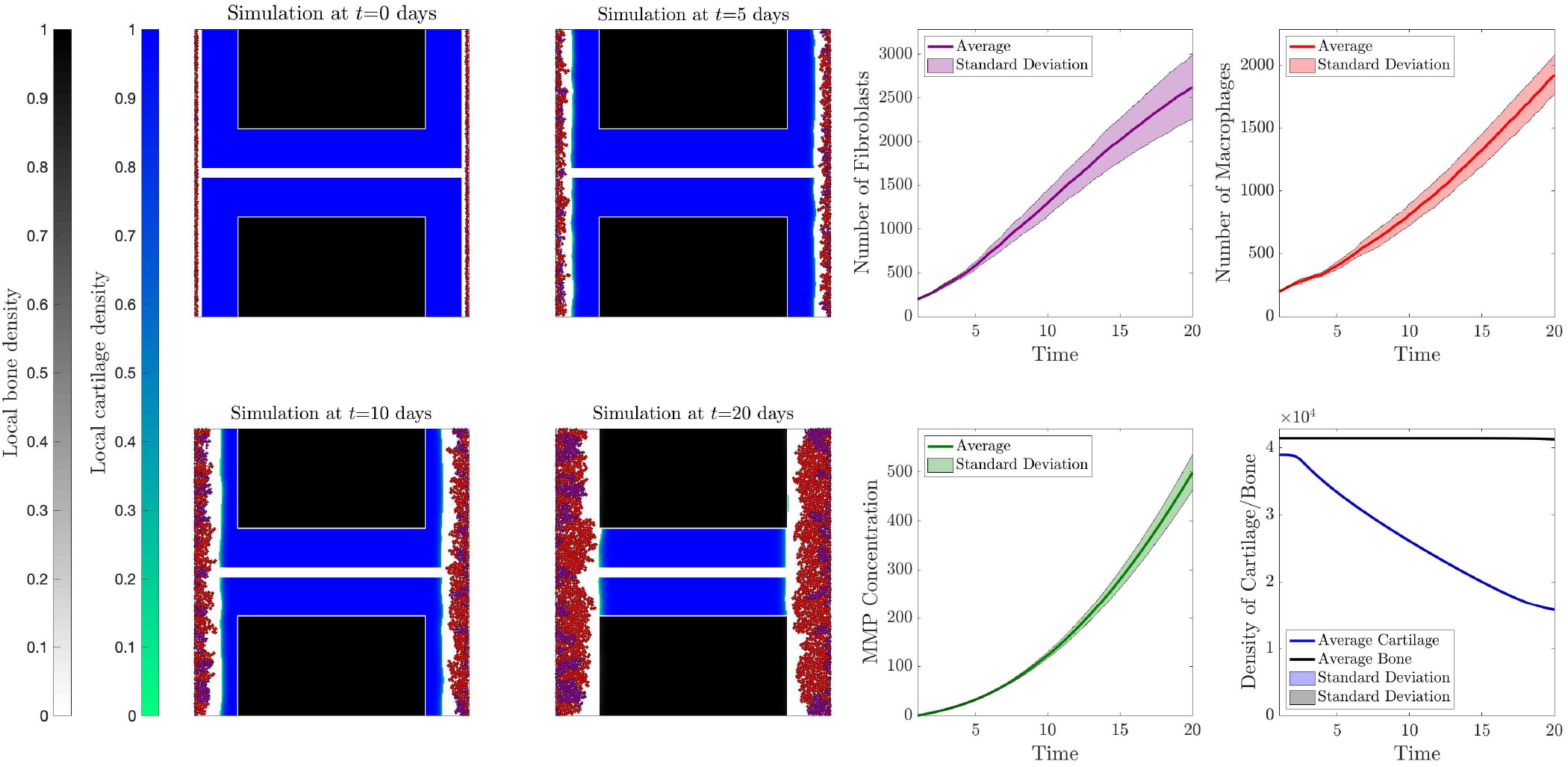
Example results where all parameters are those given in Table 2 except *α_F_* and λ*_M_* which are ten times the given value. **Left-hand plots:** Panels show the visualisation of spatial results of one of the runs of the simulation at the time-points t={0, 5, 10, 20} days. The red dots are macrophages, the purple dots are fibroblasts, the blue-green surface is cartilage density and the black-grey surface is bone density. White space represents space in which cells can move freely in the joint, *e.g*., synovial fluid. **Right-hand plots:** The plots show the cell number, concentration or density over time averaged over 5 simulation runs with the standard deviation shaded. The number of fibroblasts is given in purple, the number of macrophages in red, the global MMP concentration in green, the global cartilage density in blue and the global bone density in black.

Finally, we consider a case where we expect many more immune cells in the affected joint, to replicate an even more aggressive form of rheumatoid arthritis. We show results of the model where immune cell proliferation is increased one hundred-fold in Figure 7. Here, we observe a significantly larger increase in the number of fibroblasts and macrophages over time as expected. We also see that the concentration of MMPs is increased in comparison to the previous cases. Furthermore, the cartilage degradation is slightly more severe than the previous cases and we begin to see bone degradation occurring. Here, the standard deviation is much larger in the cell numbers. Interestingly though there is still very low standard deviation in the MMP concentration, and cartilage and bone densities. This suggests, from a modelling perspective, that even stochasticity in the number or spatial position of cells, we still expect similar levels of cartilage and bone degradation. We confirm this by plotting the final spatial distributions of each run of the simulation for the case shown in Figure 7, in Figure 8, where we observe a variety of spatial patterns arising from the cell dynamics, which do not appear to affect the overall outputs of the model.

**Figure 7:**
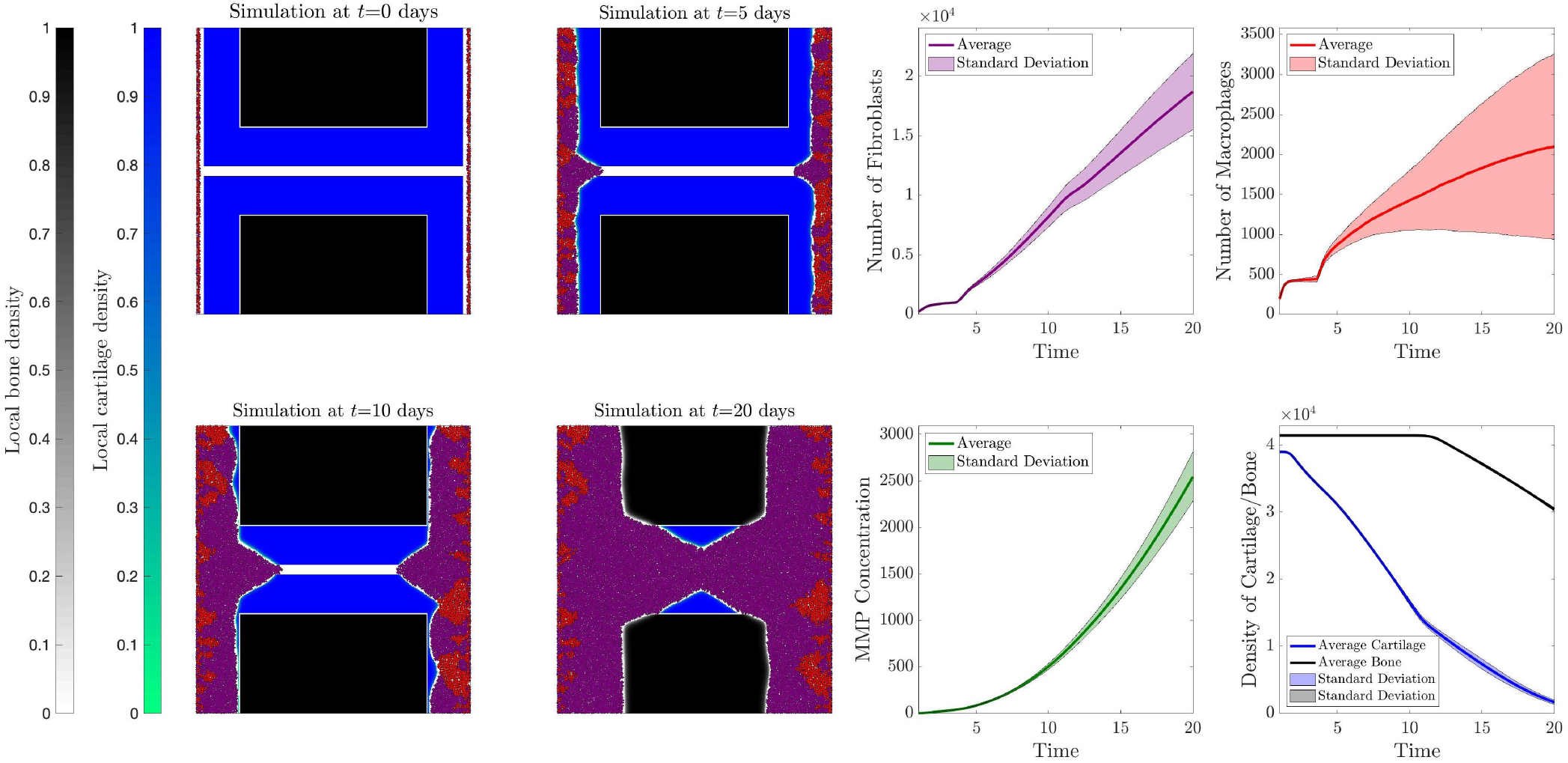
Example results where all parameters are those given in Table 2 except *α_F_* and *α_M_* which are one hundred times the given value. **Left-hand plots:** Panels show the visualisation of spatial results of one of the runs of the simulation at the time-points *t*={0, 5, 10, 20} days. The red dots are macrophages, the purple dots are fibroblasts, the blue-green surface is cartilage density and the black-grey surface is bone density. White space represents space in which cells can move freely in the joint, *e.g*., synovial fluid. **Right-hand plots:** The plots show the cell number, concentration or density over time averaged over 5 simulation runs with the standard deviation shaded. The number of fibroblasts is given in purple, the number of macrophages in red, the global MMP concentration in green, the global cartilage density in blue and the global bone density in black.

**Figure 8:**
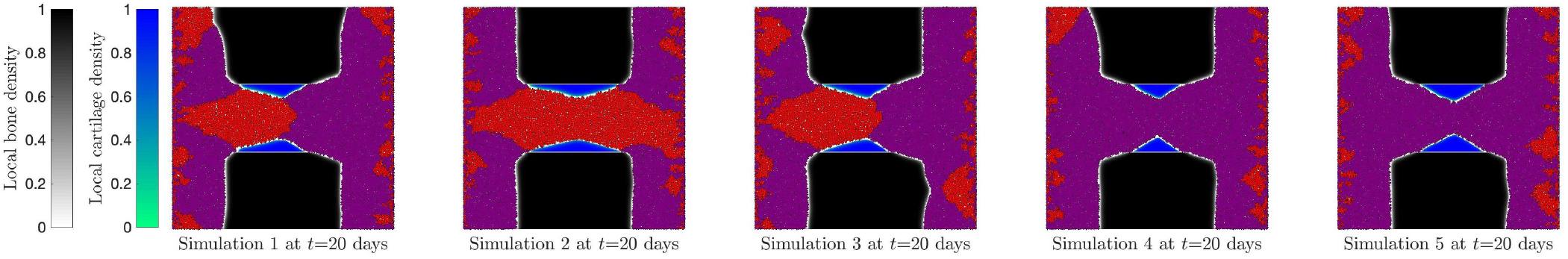
Panels show final visualisations of the 5 individual simulation runs to compare the spatial distributions of cells. The red dots are macrophages, the purple dots are fibroblasts, the blue-green surface is cartilage density and the black-grey surface is bone density. White space represents space in which cells can move freely in the joint, *e.g*., synovial fluid. Here, all parameters are base case except *α_F_* and *α_M_* which are one hundred times the base value.

### 3.3. Single parameter sensitivity analysis

The initial parameterisation of our model is based upon data from a range of theoretical and experimental work, as described in Appendix C. However, this method of parameter estimation can lead to uncertainties. Therefore, it is useful to investigate how sensitive the outputs of the model are to changes in the parameter value inputs. Sensitivity analysis techniques can help identify the key parameters within the model and identify which parameters require validated values to ensure accuracy of the model output [78, 79]. We perform a local ‘robustness’ sensitivity analysis where one single input parameter is varied while all others are kept at fixed values. We keep all parameter values to be constant, choosing the values given in Table 2 and then testing scaled values of the parameter under investigation. For each sensitivity analysis we incorporate a vector *S* to represent scalar values to multiply the parameter under investigation, where,

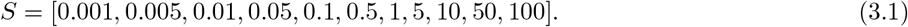

We use the notation *S_n_* = *S*(*n*) for *n* = 1, …, 11 to denote the value of each component. The maximum value of *S* is chosen to ensure all probabilities are less than or equal to 1. We then investigate parameters by setting,

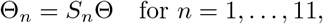

where Θ is the parameter investigated. The model outputs that we focus on are the total number of fibroblasts and macrophages over time, along with the global MMP concentration, global cartilage density and global bone density over time. Where global concentration/density refers to the sum of the concentrations/densities across all spatial positions in the domain. Each parameter setting is run 5 times and the average is plotted along with the standard deviation of these runs.

In Figure 9, we consider the movement probabilities of fibroblasts and macrophages, λ*_F_* and λ_*M*_. In the case where we vary λ*_F_* only (top row), for *S_n_* ≤ *S*_7_ = 1 there is no significant difference in all 5 outputs. However, for other values (*i.e*., *S_n_* ≥ *S*_8_ = 5) there is a gradual increase in the number of fibroblasts and global MMP concentrations as λ*_F_* increases. For larger values of *S_n_* ≥ *S*_10_ = 50 we additionally observe an increase in the degradation of cartilage, however for all *S_n_* there is no significant difference in total macrophage number or bone degradation levels. In the case where we vary λ_*M*_ only (bottom row), for *S_n_* ≤ *S*_7_ = 1 there is no significant difference in all 5 outputs. For the values of *S_n_* ≥ *S*_8_ = 5 we observe an increase in the number of macrophages, an increase in global MMP concentration and a decrease in the number of fibroblasts in the system over time as we increase λ_*M*_. Furthermore, for values *S_n_* ≥ *S*_10_ = 50 we additionally observe an increase in both cartilage and bone degradation over time. In both cases, for varying λ*_F_* or λ_*M*_, the standard deviation between runs is relatively low. These results suggest that although varying the movement probability of fibroblasts or macrophages may alter the immune cell numbers, varying these parameters does not significantly alter the cartilage and bone degradation unless large values are chosen. As each cell can only move and proliferate into free space around them increasing their movement probability allows them to find areas of free space quicker, which subsequently allows them to proliferate more freely resulting in higher numbers of cells permitted in the system, as observed.

**Figure 9:**
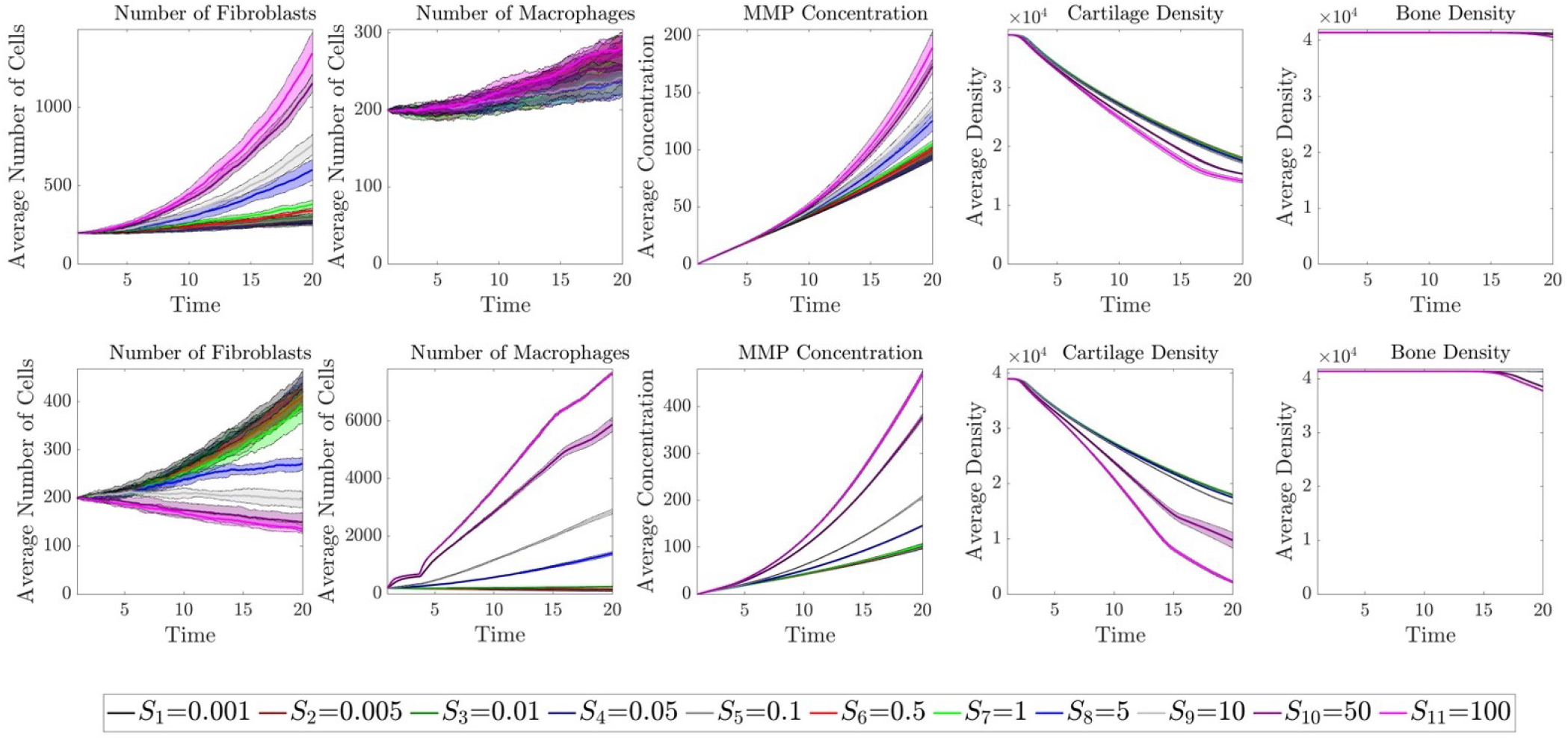
Sensitivity analyses of the movement probability of fibroblasts α_F_, (**top row**), or the movement probability of macrophages α_*M*_, (**bottom row**). The five columns represent the five key outputs of the model: the total number of fibroblasts over time, the total number of macrophages over time, the global MMP concentration over time, the global cartilage density over time and the global bone density over time. In each subplot the results for each scalar value of *S_n_* is plotted as indicated by the legend, where the original values of λ*_F_* or λ*_M_* are multiplied by *S_n_* in each case. The solid line is the average of five runs of the simulation and the shaded area in the same colour is the standard deviation between runs.

We next consider the probability of cell division for fibroblasts and macrophages, *α_F_* and *α_M_*, respectively, and each sensitivity analysis result is displayed in Figure 10. For the cases where we vary *α_F_* (top row), for the values *S_n_* ≤ *S*_7_ = 1 there is no significant difference in all 5 outputs. For the other values, *S_n_* ≥ *S_8_* = 5, we observe an increase in the total number of fibroblasts, an increase in the global MMP concentration and a decrease in the total number of macrophages as we increase *α_F_*. For the larger values of *S_n_* ≥ *S*_10_ = 50, we additionally observe a significant increase in the cartilage and bone degradation levels as we increase fibroblast cell division. Similarly, for the cases where we vary *α_M_* (bottom row), for the values *S_n_* ≤ *S*_7_ = 1 there is no significant difference in all 5 outputs. When *S_n_* ≥ *S_8_* = 5 we observe a decrease in the number of fibroblasts, an increase in the total number of macrophages and an increase in MMP global concentration as we increase *α_M_*. For the larger values of *S_n_* ≥ *S*_10_ = 50, we additionally observe a significant increase in the cartilage and bone degradation levels as we increase fibroblast cell division. Once again, in both cases, for varying *α_F_* or *α_M_*, the standard deviation between runs is relatively low. The results of these sensitivity analyses highlight that the values of *α_F_* and, to a lesser extent, *α_M_* can significantly alter the output in terms of cartilage and bone degradation. The results also highlight the competition for space between the two cell populations, where an increase in the number of fibroblasts leads to a decrease in the number of macrophages, and vice versa, in some of the parameter settings shown.

**Figure 10:**
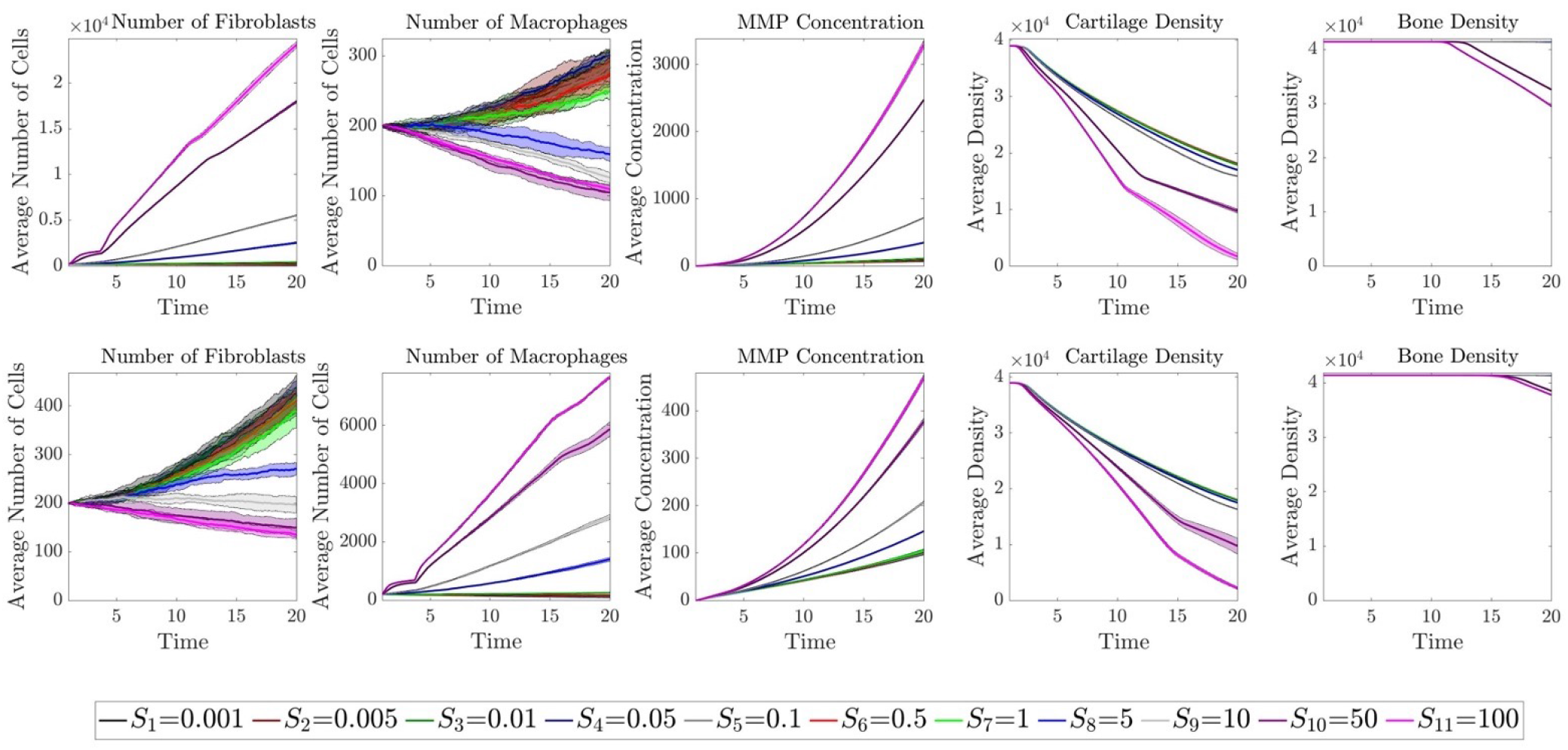
Sensitivity analyses of the division probability of fibroblasts *β_F_*, (**top row**), or the division probability of macrophages *β_M_*, (**bottom row**). The five columns represent the five key outputs of the model: the total number of fibroblasts over time, the total number of macrophages over time, the global MMP concentration over time, the global cartilage density over time and the global bone density over time. In each subplot the results for each scalar value of *S_n_* is plotted as indicated by the legend, where the original values of *α_F_* or *α_M_* are multiplied by *S_n_* in each case. The solid line is the average of five runs of the simulation and the shaded area in the same colour is the standard deviation between runs.

In Figure 11, we display the results of sensitivity analyses for varying the values of the MMP secretion rates of both fibroblasts and macrophages, *β_F_* and *β_M_*. For all values of *S_n_* the total number of fibroblasts and macrophages is consistent, as expected, as both macrophage fibroblast movement and proliferation is unaffected by *β_F_* and *β_M_*. When varying *β_F_* (top row), for the values of *S*_*n*_ ≤ *S*_7_ = 1 there is no significant change in the global MMP concentration levels, however we observe an increase in these levels once *S_n_* ≥ *S_8_* = 5. Furthermore, as we increase *β_F_*, for values *S_n_* ≥ *S_5_* = 0.1, there is an increase in the degradation of cartilage. There is no change in the bone degradation levels, until *S_n_* ≥ *S*_10_ = 50 where there is a slight increase in bone degradation. In the case where we vary *β_M_* only (bottom row), for *S_n_* ≥ 5 we observe that the global MMP concentration increases as we increase the value of *β_M_*, and the cartilage density decreases over time faster as we increase *β_F_* for values *S_n_* ≥ *S_5_* =0.1. There is additionally a small increase in bone degradation as we increase βM for values of *S_n_* ≥ *S*_10_ = 50. Once again, in both cases, for varying *β_F_* or *β_M_*, the standard deviation between runs is relatively low. The results of these sensitivity analyses highlight that the values of *β_F_* and *β_M_* can significantly alter the output in terms of cartilage degradation, but not bone degradation significantly over the time-frame considered. The results also highlight that increasing the secretion rate of fibroblasts seems to have a larger effect on cartilage degradation than increasing the secretion rate of macrophages, this could be due to the larger size of macrophages and the modelling choice that MMPs are only secreted at the centre of the cell.

**Figure 11:**
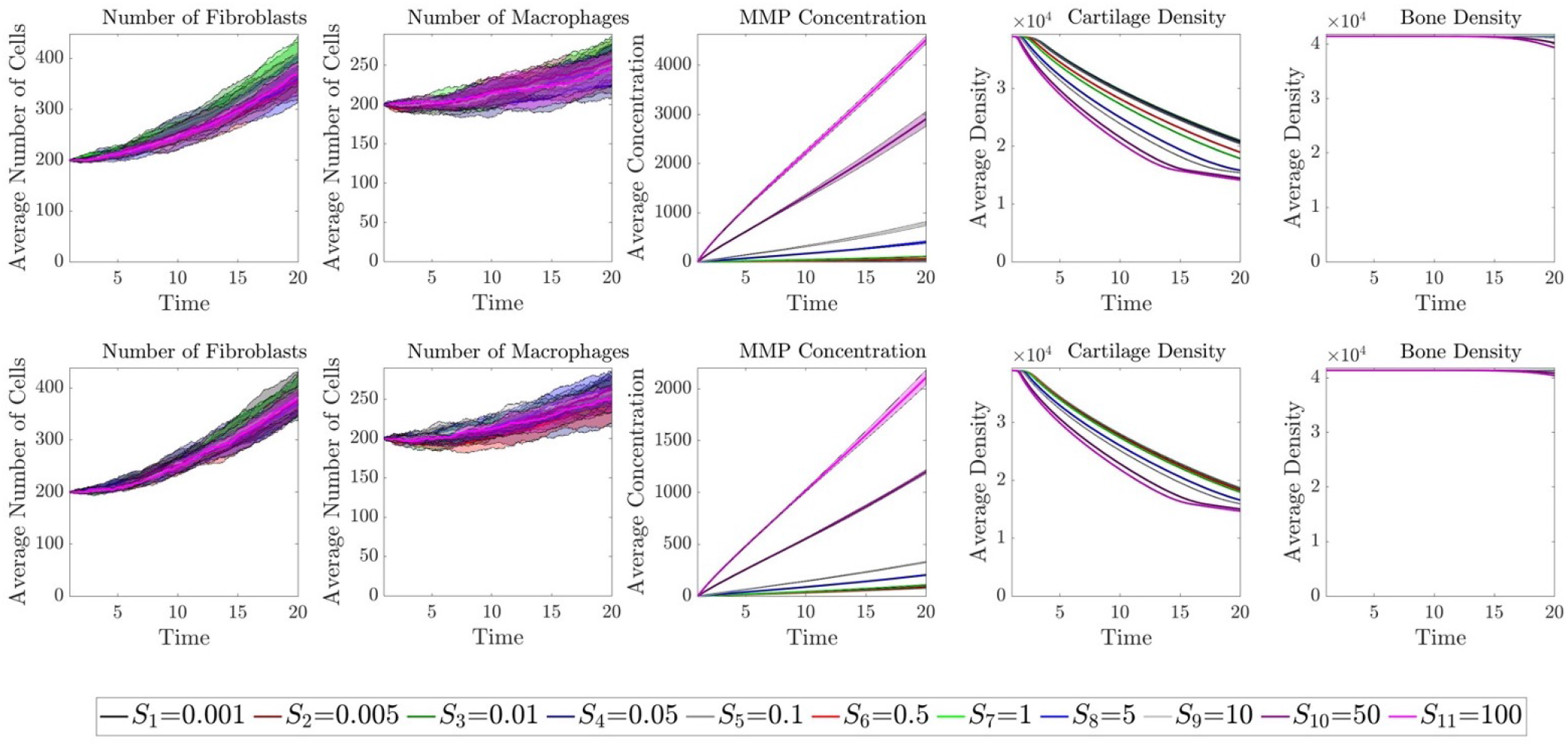
Sensitivity analyses of the MMP secretion rate of fibroblasts *κ_F_*, (**top row**), or the MMP secretion rate of macrophages *κ_M_*, (**bottom row**). The five columns represent the five key outputs of the model: the total number of fibroblasts over time, the total number of macrophages over time, the global MMP concentration over time, the global cartilage density over time and the global bone density over time. In each subplot the results for each scalar value of *S_n_* is plotted as indicated by the legend, where the original values of *β_F_* or *β_M_* are multiplied by *S_n_* in each case. The solid line is the average of five runs of the simulation and the shaded area in the same colour is the standard deviation between runs.

The sensitivity analysis results for varying the apoptosis probabilities of fibroblasts or macrophages, *κ_F_* and *κ_M_*, are displayed in Figure 12. For the case where we vary *κ_F_* only (top row), for the values of *S_n_* ≤ *S*_5_ =0.1 there is no difference in the 5 outputs and for all values of *S_n_* there is no change in the bone density levels over the time investigated. For the values of *S_n_* ≥ *S_6_* = 0.5 we observe a decrease in fibroblasts cell number and a decrease in MMP concentration levels as we increase *κ_F_*. For the values of *S_n_* ≥ *S_8_* = 5 there appears to be an increase in the total number of macrophages and a small decrease in cartilage degradation as we increase *κ_F_*. When considering the sensitivity analysis for varying *κ_M_* (bottom row), we observe that for all values of *S_n_* there is no significant difference in the global MMP concentration, cartilage density or bone density outputs. For values of *S_n_* ≥ *S_8_* = 5 there is an increase in the total number of fibroblasts as *κ_F_* increases, while for values of *S_n_* ≥ *S_6_* = 0.5 the total number of macrophages decreases as *κ_F_* increases. The results of these sensitivity analyses on *κ_F_* and *κ_M_* suggests that the values of *κ_F_* and *κ_M_* will not significantly alter the output in terms of cartilage degradation or bone degradation. However, these parameters can play a role in MMP concentration and the total number of fibroblasts and macrophages. As we increase the probability of death of one cell type, this allows the other cell type to proliferate more freely due to the increased free space, as observed in the figures displayed. As with the previous sensitivity analyses, the results are robust as we observe small standard deviation between runs for all cases.

**Figure 12:**
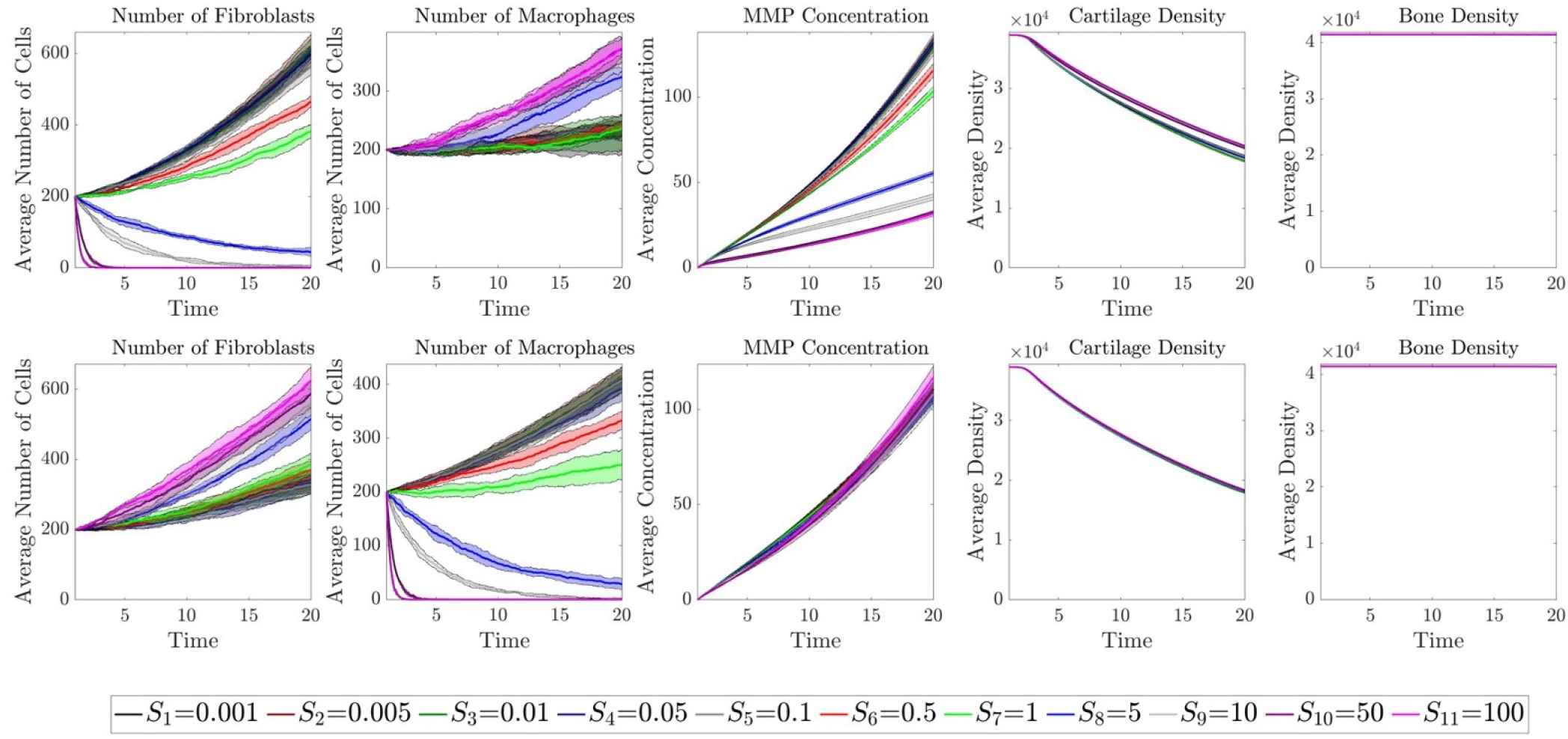
Sensitivity analyses of the apoptosis probability of fibroblasts *κ_F_*, (**top row**), or the apoptosis probability of macrophages *κ_M_*, (**bottom row**). The five columns represent the five key outputs of the model: the total number of fibroblasts over time, the total number of macrophages over time, the global MMP concentration over time, the global cartilage density over time and the global bone density over time. In each subplot the results for each scalar value of *S_n_* is plotted as indicated by the legend, where the original values of *κ_F_* or *κ_M_* are multiplied by *S_n_* in each case. The solid line is the average of five runs of the simulation and the shaded area in the same colour is the standard deviation between runs.

Next we consider the diffusion rate of MMPs, λ*_CMMP_*, and the decay rate of MMPs, *κ_CMMP_*, and perform sensitivity analyses with the results displayed in Figure 13. When varying λ*_CMMP_* (top row), for all values of *S_n_*, there is very little difference in the total number of fibroblasts, macrophages and global MMP concentrations in the model outputs. For the values of *S_n_* ≥ *S*_4_ = 0.05 there is significant increase in the cartilage degradation levels as we increase λ*_CMMP_*, while for values of *S_n_* ≥ *S_8_* = 5 we additionally observe an increase in bone degradation as λ*_CMMP_* increases. Considering the sensitivity analysis of the decay rate of MMPs, *κ_CMMP_* (bottom row), we observe that for all values of *S_n_* there is no significant change in the 5 model outputs. These results suggest that *κ_CMMP_* not a key parameter in determining the model results, while varying λ*_CMMP_* can significantly impact the output in terms of cartilage degradation or bone degradation. As we increase the diffusion rate of the MMPs, this allows the concentration in the cartilage and bone space significantly faster, promoting increased degradation. As with the previous sensitivity analyses, the results are robust as we observe small standard deviation between runs for all cases.

**Figure 13:**
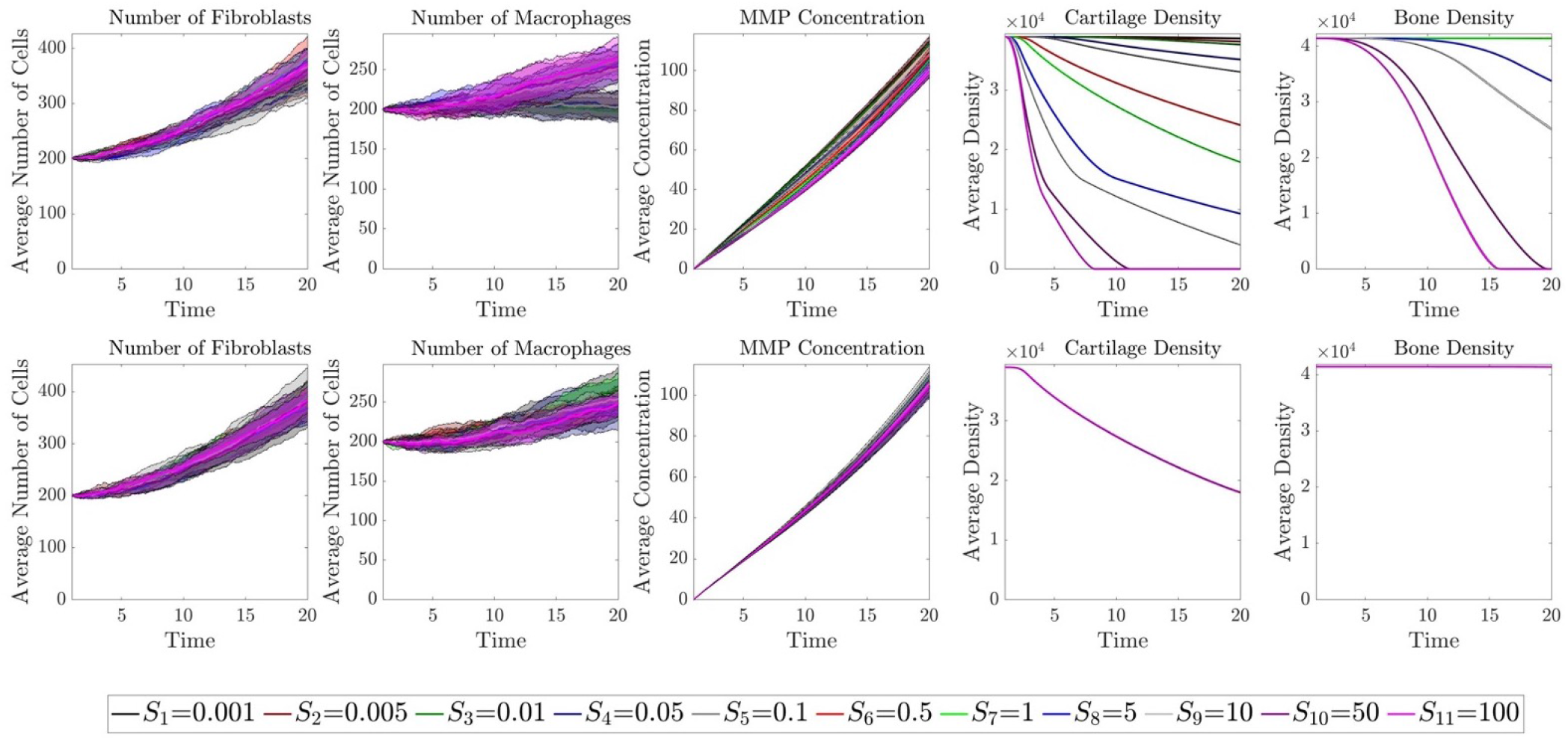
Sensitivity analyses of the diffusion rate of MMP λ*_CMMP_*, (**top row**), or the MMP decay rate *κ_CMMP_*, (**bottom row**). The five columns represent the five key outputs of the model: the total number of fibroblasts over time, the total number of macrophages over time, the global MMP concentration over time, the global cartilage density over time and the global bone density over time. In each subplot the results for each scalar value of *S_n_* is plotted as indicated by the legend, where the original value of λ*_CMMP_* is multiplied by *S_n_* in each case. The solid line is the average of five runs of the simulation and the shaded area in the same colour is the standard deviation between runs.

Finally, we perform sensitivity analyses on the degradation rates of cartilage and bone by MMPs, *κ_ρ_C__* and *κ_ρ_B__*, the results of which are displayed in Figure 14. From the results of the sensitivity analysis on *κ_ρ_C__* (top row) we observe that for all values of *S_n_* there are no significant differences in the total number of fibroblasts, the total number of macrophages, the MMP concentrations or bone density levels as the values of *κ_ρ_C__* are varied. However, for all values of *S_n_* we observe an increase in cartilage degradation as we increase *κ_ρ_C__*. From the results of the sensitivity analysis on *κ_ρB_* (bottom row) we observe that for all values of *S_n_* there are no significant differences in the total number of fibroblasts, the total number of macrophages, the MMP concentrations or cartilage density levels as the values of *κ_ρ_B__* are varied. However, for values of *S_n_* ≥ *S*_10_ = 50 we observe a very slight increase in bone degradation as we increase *κ_ρ_B__*. These results suggest that *κ_ρ_C__* and *κ_ρ_B__* only affect the output levels of cartilage and bone, respectively, and not the other outputs of the model. Furthermore, varying *κ_ρ_C__* has a larger effect on affected outputs than *κ_ρ_B__*, for the time-frame considered. As with the previous sensitivity analyses, the results are robust as we observe small standard deviation between runs for all cases.

**Figure 14:**
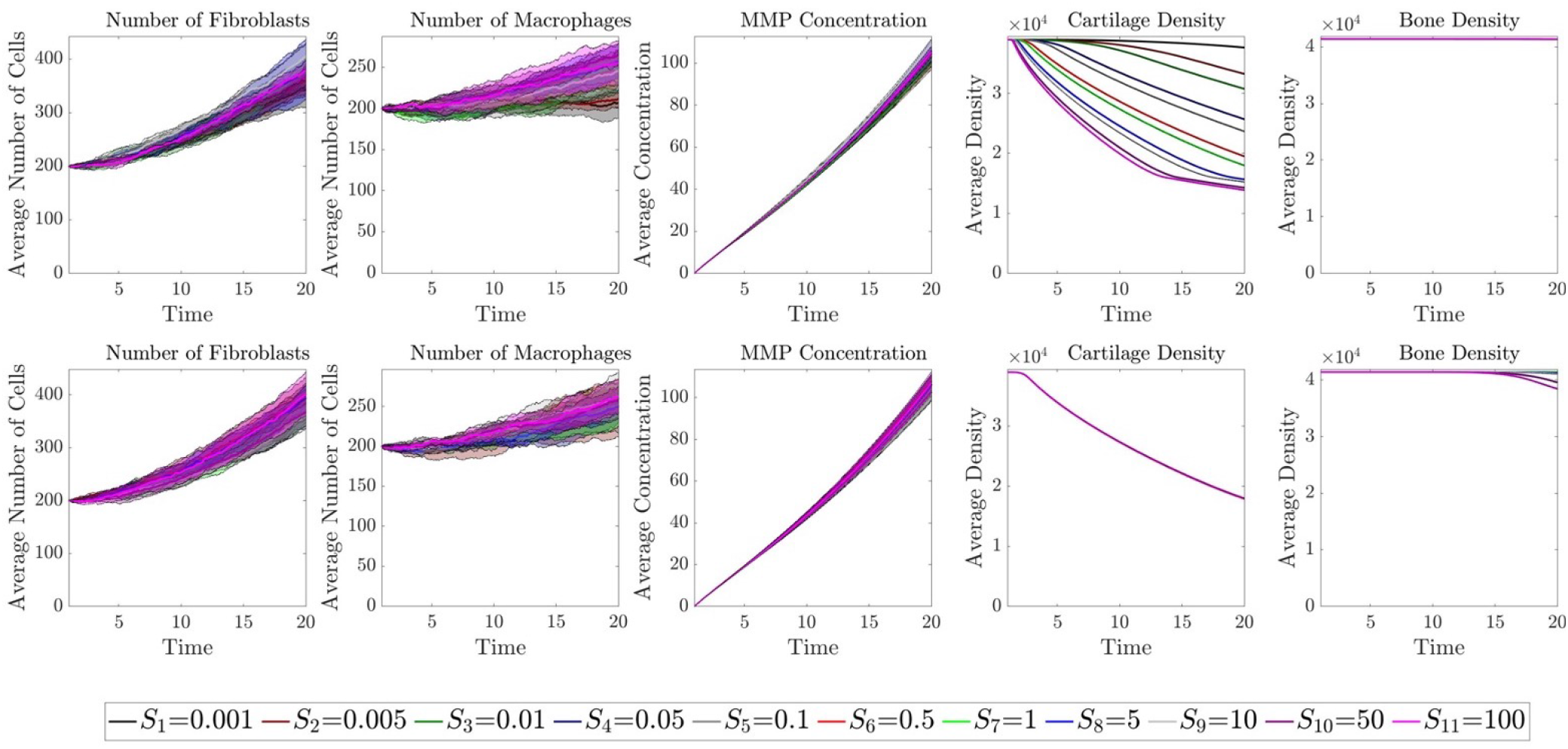
Sensitivity analyses of the MMP degradation rate of cartilage *κ_ρ_C__*, (**top row**), or the MMP degradation rate of bone *κ_ρ_B__*, (**bottom row**). The five columns represent the five key outputs of the model: the total number of fibroblasts over time, the total number of macrophages over time, the global MMP concentration over time, the global cartilage density over time and the global bone density over time. In each subplot the results for each scalar value of *S_n_* is plotted as indicated by the legend, where the original values of *κ_ρ_C__* or *κ_ρ_B__* are multiplied by *S_n_* in each case. The solid line is the average of five runs of the simulation and the shaded area in the same colour is the standard deviation between runs.

### 3.4. Relating our results to RA treatments

The modelling framework aims to describe a biological situation where no intervention (*e.g*. treatment) is included, however the results shown in Subsections 3.2 and 3.3 can be implicitly related to to several therapies currently used to treat rheumatoid arthritis. Without adding drugs explicitly to the model, we can consider the effects of treatment by varying the parameters within the model according to the mechanism targeted by the drugs. Below we discuss briefly some of commonly used treatments for RA. An overview of the drugs we could consider and the mechanisms that they promote or inhibit within the model are given in Figure 15.

**Figure 15:**
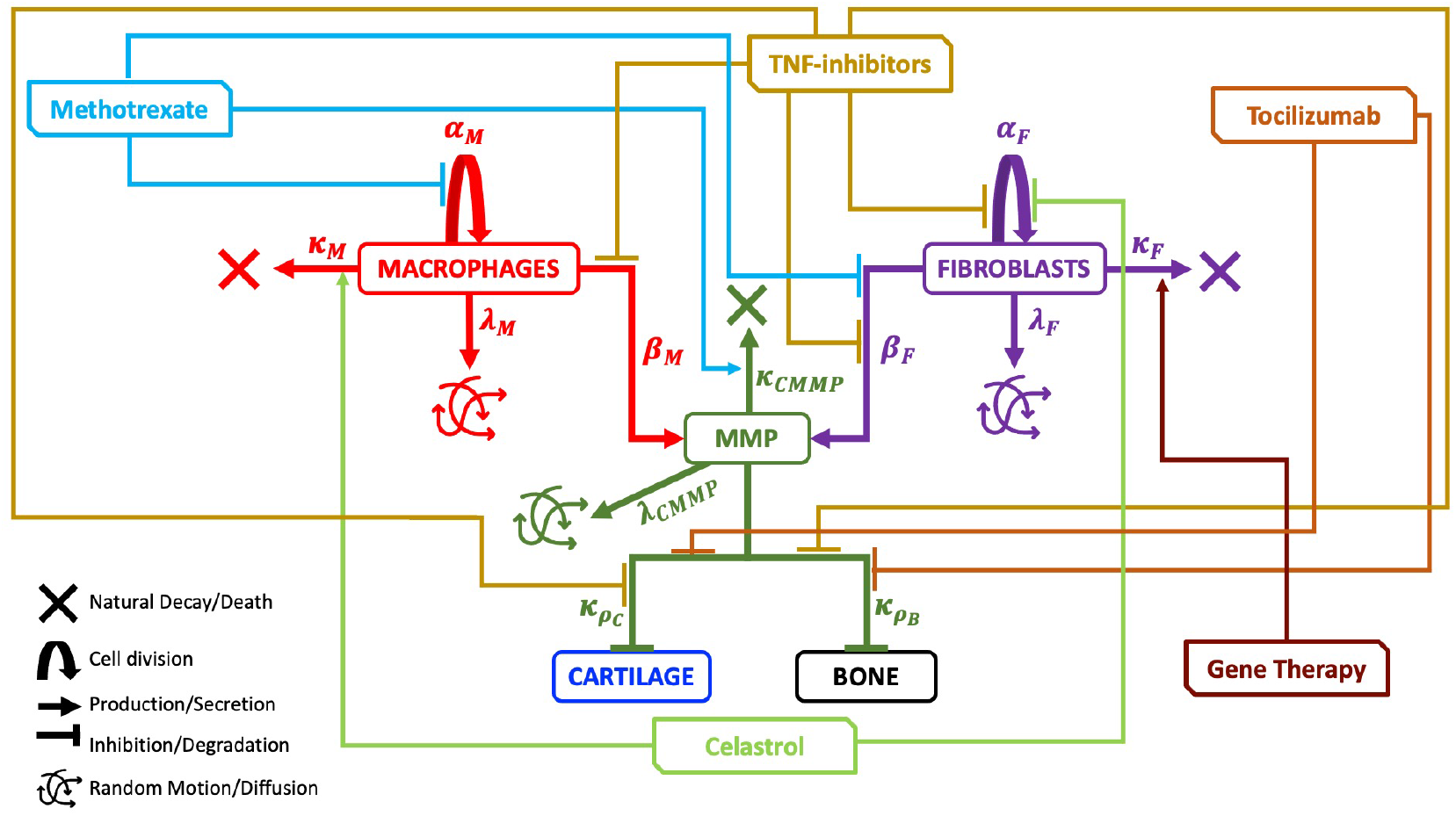
Schematic describing the mechanisms and parameters included in the hybrid model (Figure 4) with potential therapy effects included. TNF-*α* inhibitors (Gold) can inhibit fibroblast division, inhibit MMP secretion by both macrophages and fibroblasts and inhibit both cartilage and bone degradation in the RA setting. Tocilizumab (Dark orange) can inhibit both cartilage and bone degradation in the RA setting. Methotrexate (Pale Blue) can inhibit macrophage division, promote MMP decay and inhibit MMP secretion by fibroblasts in the RA setting. Gene therapy (Dark Red) can promote fibroblast apoptosis and celastrol (Pale green) can promote macrophage apoptosis and inhibit fibroblast division in the RA setting.

Disease modifying anti-rheumatic drugs (DMARDs) can target specific mechanisms within RA. For example, methotrexate, which is the most commonly prescribed drug in the UK [5, 80], has been shown to modify the cytokine profile of patients. More specifically, methotrexate can reduce IL-6 levels, which plays a role in promoting macrophage division [81], methotrexate has also been shown to inhibit secretion of MMPs by fibroblasts [5, 81] and increase the levels of TIMPs which inhibit MMPs [81]. Other RA drugs target specific cytokines within rheumatoid arthritis. Tumour necrosis factor-*α* (TNF-*α*) plays a role in promoting the inflammatory mechanisms within RA. TNF-*α* inhibitors, such as infliximab, adalimumab and etanercept are commonly prescribed DMARDs that target TNF-*α*. In untreated RA, TNF-*α* can promote the secretion of MMPs by both fibroblasts and macrophages [5], promote fibroblast cell division [5] and promote cartilage and bone degradation [5, 10]. Therefore TNF-*α* inhibitors, can reduce these effects within the RA environment. Tocilizumab is another DMARD which targets and inhibits the IL-6 receptor which is produced by several immune cell types. Among other things, IL-6 can promote cartilage and bone degradation [5, 10, 82]. By blocking IL-6 receptors, tocilizumab can reduce the effects of this cytokine. Less commonly used therapies such as chemotherapy through celastrol have promisingly been shown to reduce the aggressive proliferative abilities of fibroblasts [71] and increase apoptosis in macrophages [83]. Gene therapies can also be used to target specific cell surface receptors [84], for example, genes can be delivered that induce apoptosis in RA fibroblasts, such as with the intra-articular delivery of vectors containing PUMA, a down-stream effector of p53 and an effective inducer of apoptosis [85, 86].

The drugs considered above can be implicitly considered within our mathematical modelling framework by increasing or decreasing the parameter that controls the specified mechanism(s) targeted by the drug. We provide in Table 3 an overview of the mechanisms which each drug targets, the relevant parameter in the model and the corresponding sensitivity analysis results from Subsection 3.3 that correspond to increasing or decreasing that specific parameter without considering specific values for these parameters.

**Table 3:**
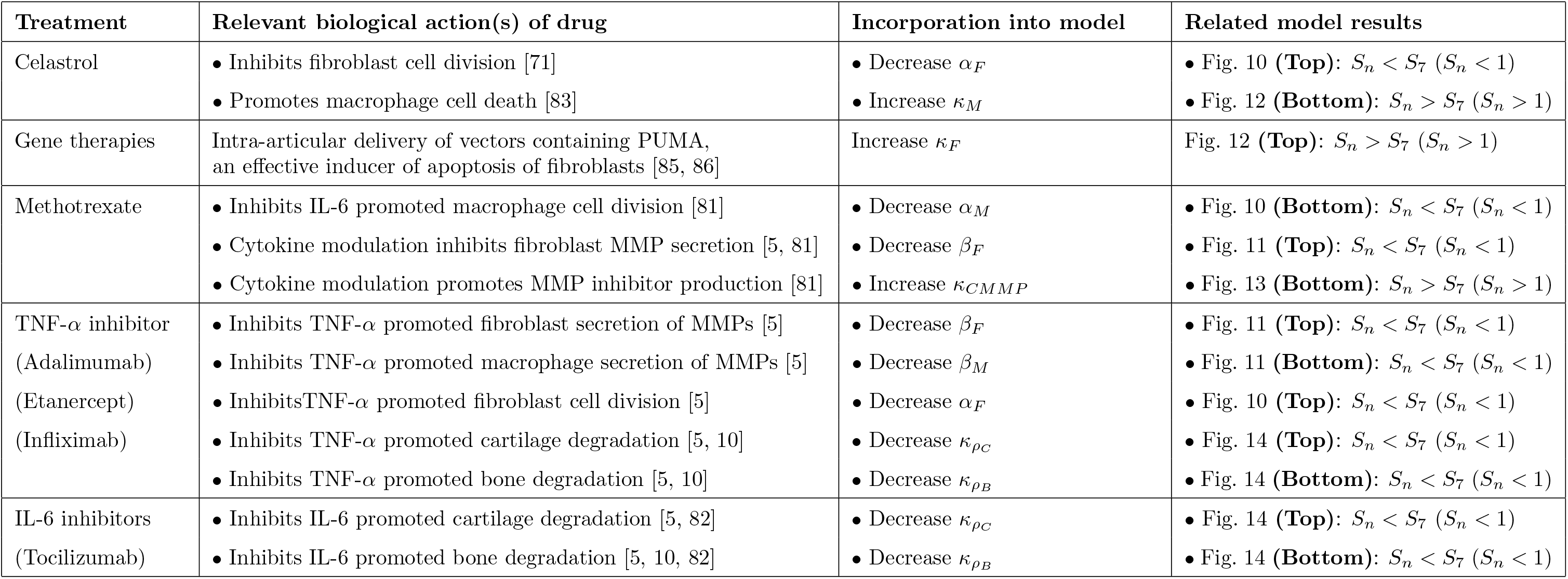
Treatments and drugs currently used to treat rheumatoid arthritis. We include the biological actions of each treatment that are relevant to the mechanisms considered within our mathematical model. We can implicitly include these mechanisms in the existing model by increasing or decreasing the relevant parameters. We provide the details of the results from Section 3.3 as an example of the effects that including these drugs implicitly could have on the model output.

Where data is available, we can consider these drugs in more specific detail within the model. For example, in [71] it was found that when celastrol is added the fibroblast proliferation rate changed from approximately 0.33 day^−1^ to 0.27 day^−1^. We can replicate this in our model by setting *α_F_* = 0.27Δ_*tcells*_ rather than the original value of *α_F_* = 0.33Δ_*tcells*_, we show the results in Figure 16. From the figure we observe that this small reduction in *α_F_* does lead to less fibroblasts in the joint, and less MMPs secreted over time. However, this change is not enough to modify the cartilage degradation levels over the time-frame considered. Therefore these results suggest that using celastrol alone, will be beneficial in reducing immune cell number, but not the overall disease outcomes in this particular parameter setting.

**Figure 16:**
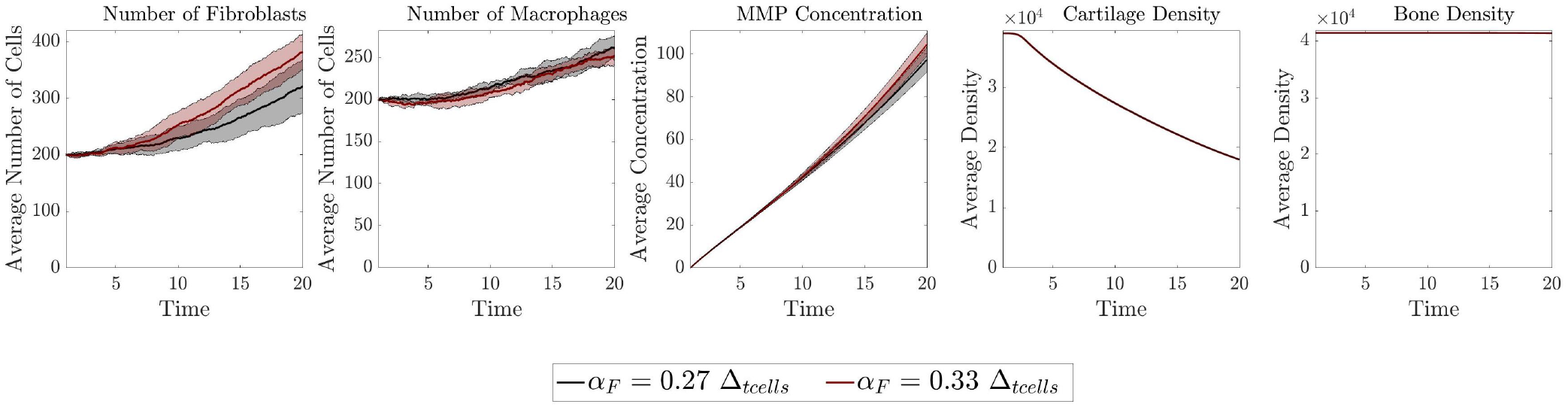
Comparing the key outputs of the model for *α_F_* = 0.27Δ*_tcells_* (black) to replicate a setting where the drug celastrol is used, compared to the original results for *α_F_* = 0.33Δ*_tcells_* (red). The five panels represent the five key outputs of the model: the total number of fibroblasts over time, the total number of macrophages over time, the global MMP concentration over time, the global cartilage density over time and the global bone density over time. The solid line is the average of five runs of the simulation and the shaded area in the same colour is the standard deviation between runs.

#### 3.4.1. Sensitivity analysis of combined characteristics

As detailed above, treatments for rheumatoid arthritis can target several mechanisms underlying the disease, rather than just a single mechanism. Therefore, it is beneficial to understand how altering more than one mechanism within the modelling framework would affect the model output. We focus on modifying two parameters at once, and as motivation for which two parameters to alter, we consider the target mechanisms of celastrol and tocilizumab.

As described above celastrol inhibits the division rate of fibroblasts [71] and promotes apoptosis of macrophages [83]. In our model this relates to decreasing the probability of fibroblast division *α_F_* and increasing the probability of macrophage cell death *κ_M_*. Therefore it is beneficial to investigate how the model outputs vary when altering both of these parameters by performing a combined sensitivity analysis, the results of which are displayed in Figure 17. The five panels represent the five key outputs of the model at the final time-step (*t* = 20): the total number of fibroblasts, the total number of macrophages, the global MMP concentration, the global cartilage density and the global bone density. In each panel the value of each output for each parameter setting is displayed as a surface plot, where low values are shown in blue increasing towards high values in yellow. Each square within the panels represents a different parameter setting. We investigate 14 values for *S_α_F__* < 1, where in the model 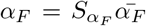 where 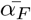 is the original value of *α_F_* given in Table 2. The values chosen are displayed below the figure as *S_*α_F_*_* (.) = *L*(.). Similarly, we investigate 17 values for *S_κ_M__* > 1, where in the model 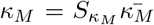 where 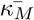 is the original value of *κ_M_* given in Table 2. The values chosen are displayed below the figure as *S_κ_M__*(.) = *M*(.). We run 5 simulations for each value of *S*_*α_F_*_ and *S_κ_M__* and average the results to understand how altering both values will alter the model outputs. From the top two panels we see that, the number of fibroblasts does not depend on the value of *κ_M_* but an increase in *α_F_* leads to an increase in the number of fibroblasts. Conversely, the number of macrophages does not depend on the value of *α_F_* but an increase in *κ_M_* leads to an decrease in the number of macrophages, in fact for values of 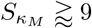 we observe extinction of the macrophage population. From the first panel in the bottom row, we find correlation between the total MMP concentration and both parameters where increasing *α_F_* leads to an increase in MMP concentration, while increasing *κ_M_* leads to a decrease in MMP concentration. For higher values of *α_F_*, we observe high concentrations of MMPs irrespective of the value of *κ_M_* suggesting that this output is more sensitive to *α_F_* than *κ_M_*. From the final two panels in the bottom row, for both cartilage density and bone density we see similar correlations for both parameters. Generally, for an increase in *α_F_* and an increase in *κ_M_* we observe larger cartilage and bone densities (less destruction). Furthermore, when *κ_M_* is high enough that we get extinction of the macrophage population, unless *α_F_* is also at the higher values we have high cartilage/bone densities (little or no destruction). The results indicate that these outputs are more sensitive to *κ_M_*. These results allow us to identify ranges for both of these parameters that result in either no change, a little change or significant change in model outputs.

**Figure 17:**
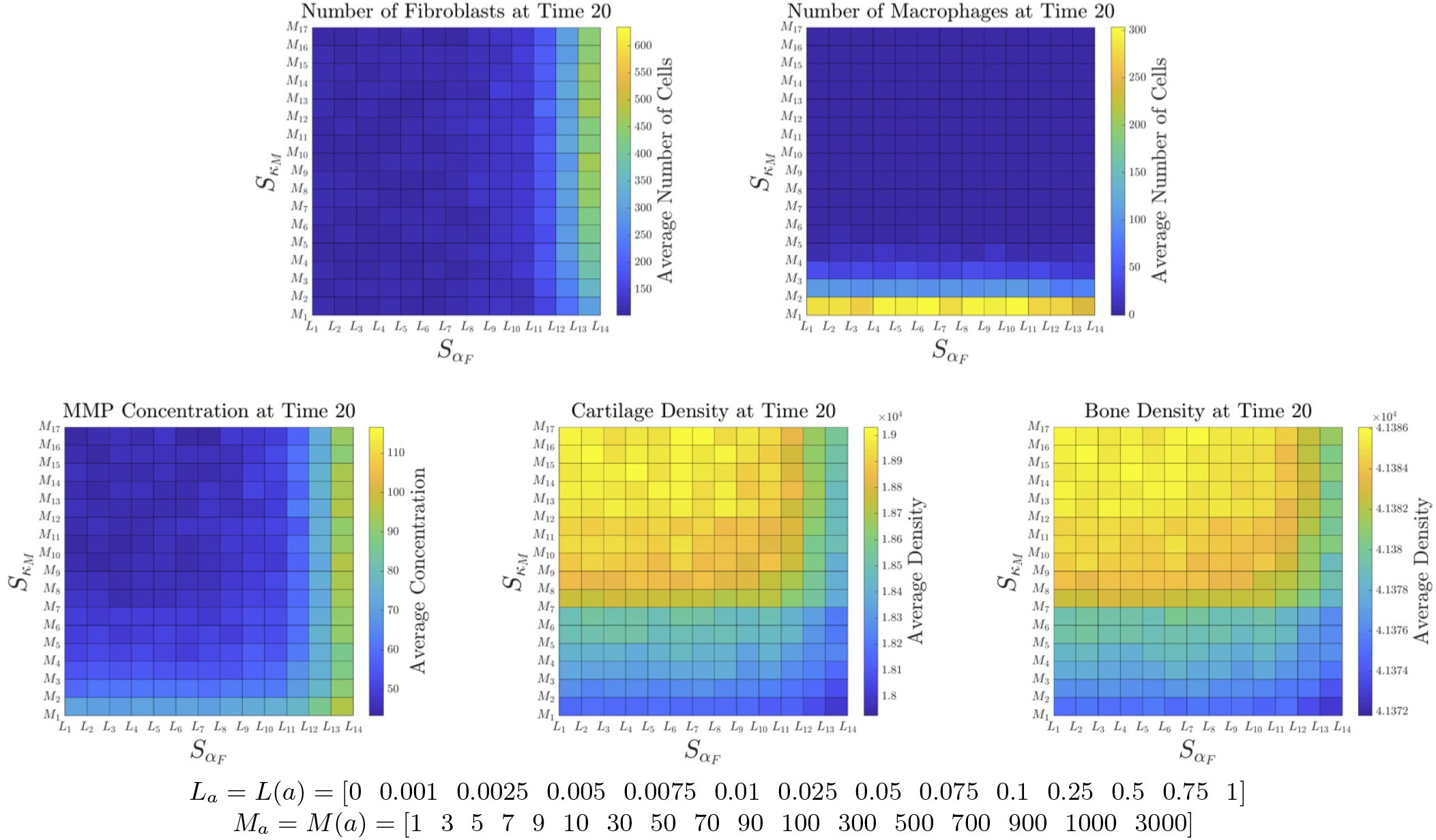
Combined sensitivity analysis of decreasing the division probability of fibroblasts (*α_F_*) and increasing the apoptosis probability of macrophages (*κ_M_*). The five panels represent the five key outputs of the model at the final time-step: the total number of fibroblasts, the total number of macrophages, the global MMP concentration, the global cartilage density and the global bone density. In each panel the averaged results for each scalar value of *S_α_F__* and *S_κ_M__* are displayed as a surface plot, where the original values of *α_F_* or *κ_M_* are multiplied by *S_α_F__* and *S_κ_M__*, respectively, in each case. The values investigated are given, where values of *S_α_F__* are taken from the vector *L*, and values of *S_κ_M__* are taken from the vector *M*. The results shown are the average of five runs of the simulation.

As a second example of investigating the outputs of the model while varying more than one parameter, we consider the mechanisms targeted by the drug tocilizumab. As described above, tocilizumab inhibits IL-6 promoted cartilage and bone degradation [5, 10, 82]. In our model this relates to decreasing both the decay rate of cartilage by MMPs, *κ_ρ_C__*, and the decay rate of bone by MMPs, *κ_ρ_B__*. To investigate how the model outputs vary when altering both of these parameters we consider combined sensitivity analysis, the results of which are displayed in Figure 18. The five panels represent the five key outputs of the model at the final time-step (*t* = 20): the total number of fibroblasts, the total number of macrophages, the global MMP concentration, the global cartilage density and the global bone density. In each panel the value of each output for each parameter setting is displayed as a surface plot, where low values are shown in blue increasing towards high values in yellow. Each square within the panels represents a different parameter setting. We investigate here, 8 values for both 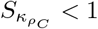 and 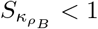, that is decreasing values for both parameters. The simulations are run such that 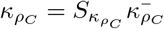 and 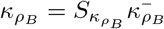 where 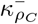 and 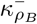 are the original decay rates given in Table 2. The values investigates are displayed below the figure as 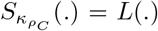 and 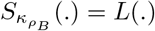. The results suggest that, generally, there is no significant difference in the number of fibroblasts, number of macrophages or the concentration of MMPs when varying *κ_ρ_C__* and *κ_ρ_B__*, as the small differences in these panels can be attributed to the stochasticity within the model. From the final two panels in the bottom row, for both cartilage density and bone density we see that each output only depends on one of the parameters. That is, increasing *κ_ρ_C__* leads to an increase in cartilage destruction, irrespective of the value of *κ_ρ_B__*, and conversely *κ_ρ_B__* leads to an increase in bone destruction, irrespective of the value of *κ_ρ_C__*. These results further highlight that there is no difference in model outputs when considering variations in both *κ_ρ_C__* and *κ_ρ_B__*, compared to varying only one of these parameters. Therefore, for the mechanisms targeted by tocilizumab, in the model, there is no correlation between the decay rate of cartilage and of bone by MMPs (Figure 18) while such a correlation seems to exist for the mechanisms targeted by celastrol (*i.e*., fibroblast division and macrophage cell death, Figure 17). Since experimental studies have shown that tocilizumab seems to affect both degradation of bone and cartilage, this suggests that there could be other detailed biological mechanisms for tocilizumab that were not considered in this study, and should be added to allow for more realistic descriptions of this drug.

**Figure 18:**
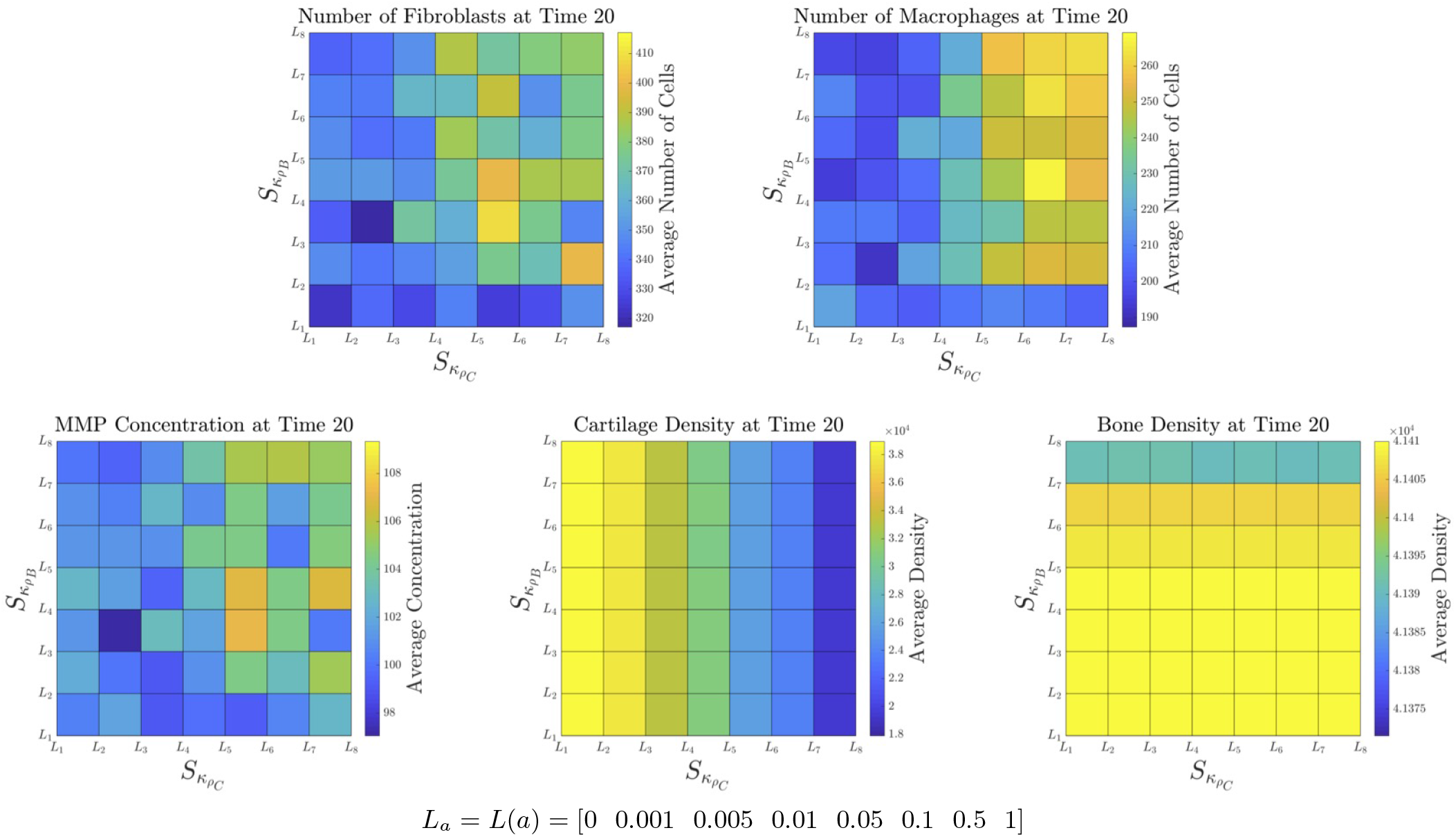
Combined sensitivity analysis of decreasing the decay rate of cartilage by MMPs (*κ_ρ_c__*) and decreasing the decay rate of bone by MMPs (*κ_ρ_B__*). The five panels represent the five key outputs of the model at the final time-step: the total number of fibroblasts, the total number of macrophages, the global MMP concentration, the global cartilage density and the global bone density. In each panel the averaged results for each scalar value of 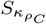 and 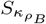 are displayed as a surface plot, where the original values of *κ_ρ_C__* or *κ_ρ_B__* are multiplied by 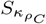 and 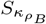, respectively, in each case. The values investigated are given, where values of 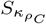 and 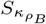 are both taken from the vector L. The results shown are the average of five runs of the simulation.

## 4. Discussion and conclusions

As a first step towards building a more realistic model of arthritic destruction within a small PIP joint, in this study we introduced an off-lattice hybrid model to describe the growth and development of the pannus in the PIP joint space, in the context of rheumatoid arthritis (RA). The model combined an off-lattice individual-based approach for the cell dynamics coupled with a deterministic PDE approach to describe the dynamics of MMPs, cartilage and bone. As previously highlighted, this model is an initial formulation and a first step towards building a more realistic model of pannus formation within a rheumatoid arthritis affected joint. Therefore at this stage the model cannot be used to predict progression of disease accurately. However, the model can be used to understand the effects that altering the mechanisms included on the simulation results, highlighting the key parameters that require further validation.

We have provided some initial results that considered three scenarios with increasing proliferation rates of cells to investigate increasing aggressiveness of the pannus progression (Figures 5–7). The model results suggest that increasing the proliferation rates of cells ten-fold does not lead to a significant increase in the destruction of bone, even with significantly increased numbers of cells (comparing Figures 5 and 6), while increasing the proliferation rates of cells a hundred-fold did result in increased levels of bone destruction (comparing Figures 5 and 7). This suggests that the latter parameter setting can be used to describe a more aggressive form of the disease. Furthermore, even when we observed spatial heterogeneity between simulation results, we still obtain low standard deviation in the global MMP concentrations, global cartilage density and global bone density outputs (Figure 8), indicating that varied spatial patterns in the pannus, can still result in the same outcomes and rate of degradation in the joint. We note that, this initial model is not yet at a stage where direct comparisons to clinical and experimental data can be made. However, some level of biological insight can be obtained through considering the level of variation in the model outputs when increasing or decreasing the effects of the biological mechanisms included (*i.e*., varying the model parameters). Moreover, extensions of the model to contain appropriate levels of complexity could be validated with more accurate data to allow for direct comparison with the spatial patterns of cell populations within joints affected by RA, such as those seen within studies that image the progression of RA [87–91].

To identify the role of the key parameters within the model we performed a one-at-a-time sensitivity analysis. These sensitivity tests allow us to identify the parameters that, when varied, will significantly alter the outputs of the model over the time-frame considered. The results highlighted that varying the immune cell movement probabilities (Figure 9), varying the apoptosis rates of cells (Figure 12), varying the decay rate of MMPs (Figure 13) or varying the degradation rate of bone by MMPs (Figure 14) did not significantly affect cartilage and bone degradation over the time-frame considered compared to the variation of other parameters. On the other hand, the results also showed that bone and cartilage density can be significantly reduced when we increase the division probability of cells (Figure 10), the MMP secretion rates of cell (Figure 11), the diffusion rate of MMPs (Figure 13) or the degradation rates of cartilage (Figure 14), when compared to the variation of other parameters. Therefore, the outputs of the model are more sensitive to these parameters and for the model to be able to provide biologically relevant results we require the values of these specific parameters to be as accurate as possible. To obtain accurate values for these parameters, data from experimental and clinical studies are required.

In Section 3.4, we discussed the above sensitivity analysis results in the context of current therapeutic approaches for RA. In particular, we emphasised the model parameters that could be varied to investigate (implicitly) the effects of different drug therapies, and how these changes in model parameters could be inferred by our sensitivity analysis numerical simulations results. Furthermore, focusing on the mechanisms targeted by celastrol and tocilizumab for motivation, we briefly investigate combined sensitivity analyses to investigate the effects of altering more than one parameter on model output(s). We can infer the general changes to model output we expect with increasing or decreasing the parameters that can be affected by the drug, highlighting the expected qualitative effects of these drugs on the key outputs of the model. We note that the specific inclusion of the dynamics of these drugs into the mathematical model, which was not the purpose of this particular study, could lead to more complex dynamics due to potential nonlinear effects between different system components. This will be the focus of a future study.

### 4.1. Future work

As described in Section 1.1, there are various layers of biological complexity that have not been included in the model so far. This simple framework can be easily extended using the current methods to incorporate further biological complexity. We provide some suggested extensions to the model here.

The current model considers only resident cells within the joint space, however immune cells can also be recruited from the circulatory system and promote angiogenesis promoting this recruitment [5, 13, 14]. We can extend the model to allow new immune cells to enter the domain at the boundaries, or more specifically introduce vasculature a the boundaries of the domain and allow the cells to enter through blood vessels. Moreover, in RA, the growth of the pannus can result in local hypoxia driving angiogenesis which can further promote an influx of pro-inflammatory immune cells entering the joint [14]. If we include vasculature we can also consider the influx of oxygen and nutrients into the system, modelling these chemicals in as similar way to the MMPs in this work. The recruitment of cells through vasculature could be incorporated using methods similar to individual-based approaches used to model the metastasis of tumour cells through vasculature [92].

Currently we focus on homogeneous populations of immune cells, where all cells within the population exhibit the same probabilities of migration and proliferation. However, the phenotype of fibroblasts can be altered to a more tumour-like state where cell division is increased, apoptosis of cells is decreased and migration of cells is more invasive [16, 87, 93, 94]. There can be similar heterogeneity within the macrophage populations [15]. We could incorporate this within the model by tracking the phenotype of each fibroblast and macrophage and allow the cells to alter their phenotypes over time. In the model we consider only resident fibroblasts and resident macrophages, and not subsets of these populations or other immune cell types. Further immune cells can play a role in the progression of rheumatoid arthritis such as T cells and B cells [13, 95]. These immune cells, their mechanisms and their interactions with components of the joint could be included within the model using the same methods as those used to incorporate fibroblasts and macrophages.

As mentioned in Section 2.3, we implicitly take into account the effects of tissue inhibitors of MMPs (TIMPs) by including decay of MMPs and limitations on the local concentration of MMPs. We could take explicitly include TIMPs, using PDEs and further investigating the activator-inhibitor like mechanisms of MMPs with TIMPs. To do this we could utilise a system of equations similar to that used in [39] to model MMP and TIMP dynamics.

Cartilage and bone densities are considered at a tissue-level scale within the framework. However we could explicitly incorporate the cellular-level components of these tissues. For example, the role of chondrocytes and the highly organised extracellular matrix (ECM) within cartilage [3, 5, 10, 13] could be considered. Furthermore, the (im)balance between osteoclasts, which degrade bone, and osteoblasts, which produce bone, within the bone [5, 10, 13] could also be investigated in the RA context further. To incorporate these features we could consider method similar to those used to previously model cartilage growth [96] or bone remodelling [97–102].

All of the above mechanisms that can be incorporated into the model would extend the complexity of the model and the number of parameters required. Therefore, to incorporate this biological complexity and to obtain biological relevance, we would require recent clinical and experimental data to validate the modelling choices and parameter settings investigated.

## Acknowledgments

This research did not receive any specific grant from funding agencies in the public, commercial, or not-for-profit sectors.

## Conflict of interest

All authors declare no conflicts of interest in this paper.

## Appendix A. Discretisation of PDEs

We require an on-lattice approach for numerical simulations of the PDE components of the model, to perform simulation we discretise the PDEs in space and time to allow for this. We consider the lattice to have spatial index positions *i* ∈ [1, …, *N_x_*] and *j* ∈ [1, …, *N_y_*], and consider the time to be discretised so that the time-step *k* = *t*/Δ*_tchem_* where Δ_*tchem*_ is some time-step length. We can then discretise the PDE in Equation (2.1) using a fully explicit form to be written as,

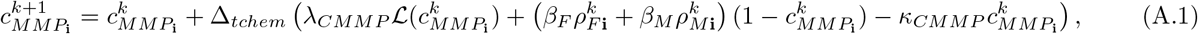

where 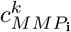 is the concentration of MMP at spatial position **i** at time-step *k*, Δ*_tchem_* is the time-step length and 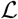 is the finite difference Laplacian, such that,

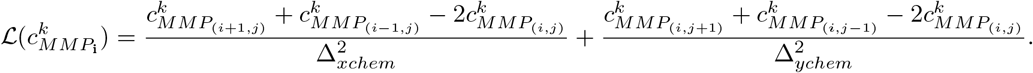

Here, Δ*_xchem_* and Δ*_ychem_* are the space-step lengths in the *x* and *y* direction, respectively. Similarily, the discrete forms of bone and cartilage equations Equations (2.2) and (2.3), can be written as,

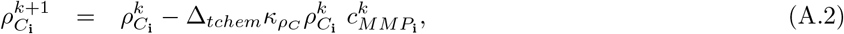

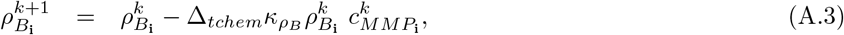

where 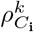 and 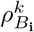 are the densities of cartilage and bone at spatial position **i** at time-step *k*, and Δ*_tchem_* is the time-step length.

## Appendix B. Code availability

The full code used to perform the numerical simulations in Matlab is available upon request via email to the corresponding author or can be accessed on GitHub at: https://github.com/frm3-st-andrews/Arthritis.

## Appendix C. Parameter estimation

### Appendix C.1. Time and space-steps of numerical simulations

For the off-lattice description of cells we arbitrarily choose the time-step of simulations to be,

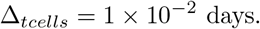

For the discretised PDEs describing MMPs, cartilage and bone we impose a time-step Δ*_tchem_* and space-steps in the *x* and *y* spatial directions, Δ*_x, y_*, for the numerical simulations such that the finite difference method [103] used to solve the equations are stable. We choose,

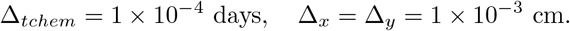

Note that, we choose a different time-step for the deterministic part of the model than the stochastic. This choice is made to speed up the simulations where possible, as a smaller time-step is not required for the stochastic parts.

### Appendix C.2. Set-up of the spatial domain

We define the full domain of the grid to be *x* ∈ [*x_l_, x_h_*] and *y* ∈ [*y_l_*,*y_h_*]. We consider the length of the domain in the y direction depends on the choices of the sub-domain sizes described in Table 1. Therefore we set,

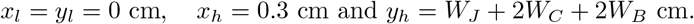

We arbitrarily choose the height of bone in the *y* direction protruding into the top and bottom of the domain to be,

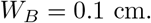

In [7], the authors use high resolution MRI (magnetic resonance imaging) to evaluate the cartilage of the 2nd and 3rd, metacarpophalangeal (MCP) and proximal interphalangeal (PIP) joints in hand osteoarthritis patients and healthy controls. Based in the Netherlands, 41 OA patients and 18 healthy controls were evaluated. The patients were all female, the OA patients were between 40-80 years old, while the healthy individuals were between 18-31 years old. For healthy controls the cartilage thickness in the PIP joint varied between 0.2mm and 0.7mm, with a mean of 0.4mm and standard deviation of 0.1mm. This mean and standard deviation was the same with the OA patients, but they exhibited higher levels of variation. Therefore, we consider that prior to pannus formation, the cartilage width in our simulated PIP joint will be approximately 0.4mm, as this was the average of healthy controls. Therefore, we consider the cartilage to be uniform width surrounding the bone and set,

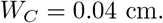

In [6], the authors use radiographs to assess the joint space width (JSW) in undamaged MCP and PIP joints of patients with early rheumatoid arthritis (RA). The mean JSW of the 4 fingers on both hands was measured for the each of the 38 patients that were assessed. The cohort was made up of 29 females and 9 males, 13 patients were under 50 years old, 8 patients were 50-60 years old, and 17 were over 60 years old. In the males the average joint space width was 1.28 mm (±0.1mm for the 95% confidence interval), while females were 0.99mm (±0.05mm). The averages between the age groups were all roughly the same, (1.1mm, 1.06mm, 1.02mm, respectively). Similarly, in [8] the authors use radiographs to validate the joint space width (JSW) calculations in undamaged MCP and PIP joints of patients with early rheumatoid arthritis (RA). Two dutch data sets were used to validate the method, where radiographs were taken every 6 months of the patients. They authors use data from the 4 fingers on each hand, of each patient (omitting the thumbs). For the 1st data set, the average PIP joint in undamaged cases (527 images tested) was 1±0.2 mm, while in the 2nd data set (570 images tested) the average was 0.9±0.2mm. Therefore, we consider that prior to pannus formation, in our simulated PIP joint we would expect the initial joint width to be approximated as,

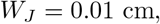

where we assume the spacing to be uniform across the width of the joint. In health, the synovial membrane contains relatively few cells consisting of an intimal layer of 1-2 cell thickness and a distinct sublining [11]. Therefore we initially want to allow around 2 layers of cells in the membrane, so we choose

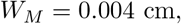

This value is set so that an approximate maximum thickness of two macrophages is possible for the cell membrane. We arbitrarily then choose the space between the cartilage and the edge of the domain in the x-direction to be,

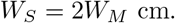

### Appendix C.3. Cell parameters

#### Cell radii

In [76], the authors show that the average diameter of human alveolar macrophages from the lung of 10 participants was 21.2±0.3 *μ*m in diameter. They also highlight that the diameter of human macrophages is much large than that of other species. Furthermore, in [75] it was shown that the average diameter of a fibroblast is between 10-15*μ*m, while for a macrophage the average diameter can be as large as 20-80*μ*m. Therefore, we choose the diameter of a macrophage to be approximately 21*μ*m and the diameter of a fibroblast to be approximately 13*μ*m, that is we set,

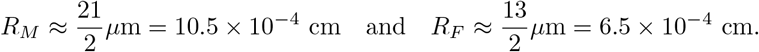

#### Initial condition

In a healthy joint, the intimal layer of the membrane is generally 1-2 cells thick and consists of fibroblast-like synoviocytes (FLSs) and macrophage-like synoviocytes (MLSs) evenly distributed and in equal amounts [11, 12]. Therefore we set the initial number of both cell types to be equal as,

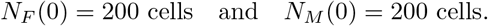

#### Cell division

In [71], the authors study the effects of celastrol, and anti-inflammatory chemical, on the proliferation rates of fibroblast-like synoviocytes from RA patients. In the untreated case the approximate doubling time of FLSs is 2.1 days. This can be used to estimate an approximate growth rate through the equation for exponential growth, 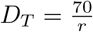, with *D_T_* being the doubling time and *r* the % growth rate. Using this we can estimate the FLSs will have a proliferation rate of approximately 0.33 day^−1^, which corresponds to the probability of a fibroblast dividing at any time-step as,

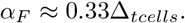

The authors in [39] also use a similar value for fibroblast division and estimate that macrophage division will be similar. Therefore, we use in our simulations that,

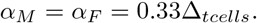

Note, that these probabilities are within the context of the cell having available space. As macrophages are larger than than fibroblasts, they are less likely to find available space to divide within the simulations, and therefore the probability of a macrophage dividing will be inherently lower than that of fibroblasts. This is consistent with the suggestion that while the proliferation of both macrophages and fibroblasts within the pannus increases, fibroblasts may exhibit a particularly aggressive phenotype and have higher proliferation rates [87, 93, 94].

#### Cell death

In [72], the authors use experimental methods to detect apoptotic cell death in synovial tissues biopsied from 6 patients with RA, and 3 with OA as controls. They found that 30% of the RA fibroblast like-synoviocytes were susceptible to apoptosis. The authors of [39], use this value and the assumption that apoptosis takes 24hrs to estimate that the apoptosis rate of fibroblast like-synoviocytes is around 0.3 day^−1^. This would lead to the probability of a fibroblast undergoing apoptosis to be,

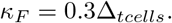

This value results in proliferation and decay being very close (similar to a homeostatic situation) but leads to extinction due to stochasticity. At the moment we choose a value of 10% of this, to prevent fibroblast extinction and assume the value would be similar to macrophage apoptosis rates. In the review paper [73], the authors discuss the key phenotypes of monocytes and macrophages. Specifically, they discuss mouse model results, where Ly6C+ intestinal monocytes that have a half-life of 3 weeks. The authors of [39], use this mouse model value and assume that apoptosis of macrophages occurs at the rate of 0.033 day^−1^. Therefore, we set the probability of a macrophage undergoing apoptosis to be,

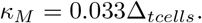

#### Cell migration

In many mathematical modelling works animal cell movement is estimated to be of order 1 × 10^−11^ cm^2^ sec^−1^ [39, 74, 104–106]. This estimate translates to a diffusion rate of 8.64 × 10^−7^cm^2^ day^−1^, which through scaling corresponds to a probability of moving of 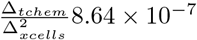. Therefore, we set for fibroblasts,

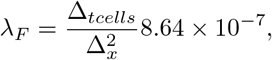

and similarly for macrophages,

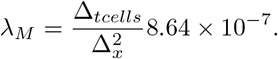

We note that these are just estimates and, as macrophages are larger, they may have a lower movement probability. Also note that λ is not the total probability, but contributes to this probability as we also consider a volume exclusion process whereby cells cannot overlap in space with each other or with cartilage or bone. This probability relies on Δ*_tchem_* and Δ*_xcells_* = Δ*_x_* which have to be chosen such that λ ≤ 1.

### Appendix C.4. Parameters for MMPs, cartilage and bone

#### MMP secretion rates of cells

In the mathematical model described in [39], the authors estimate the MMP secretion rates of both fibroblasts and macrophages. For our initial simulations we use these values and set,

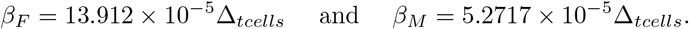

#### Diffusion and decay of MMPs

The parameters for the mechanisms of MMPs in the joint are taken to be those used in [39]. They estimate the diffusion rate of MMPs, using [74], to be,

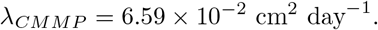

Furthermore, they estimate, using [77], the natural decay rate of MMPs to be,

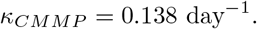

#### Degradation of cartilage and bone by MMPs

In [39], they only consider cartilage degradation, which they estimate to occur at the rate,

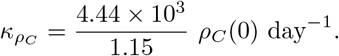

We make the assumption that the decay rate of bone will be lower, and for simulations, take the value,

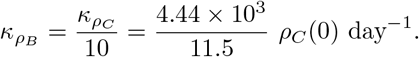

## Notes

https://github.com/frm3-st-andrews/Arthritis

### Competing Interest Statement

The authors have declared no competing interest.

### Summary of Updates

The manuscript has been revised to make the modelling approach clearer and highlight the benefits of the modelling framework

